# Motif-VI Loop Acts as a Nucleotide Valve in the West Nile Virus NS3 Helicase

**DOI:** 10.1101/2023.11.30.569434

**Authors:** Priti Roy, Zachary Walter, Lauren Berish, Holly Ramage, Martin McCullagh

## Abstract

The flavivirus NS3 helicase (NS3h), a highly conserved protein, plays a pivotal role in virus replication and thus represents a potential drug target for flavivirus pathogenesis. NS3h utilizes nucleotide triphosphate, such as ATP, for hydrolysis energy (ATPase) to translocate on single-stranded nucleic acids, which is an important step in the unwinding of double-stranded nucleic acids. The intermediate states along the ATP binding and hydrolysis cycle, as well as the conformational changes between these states, represent important yet difficult-to-identify targets for potential inhibitors. We use extensive molecular dynamics simulations of apo, ATP, ADP+P_i_, and ADP bound to WNV NS3h+ssRNA to model the conformational ensembles along this cycle. Energetic and structural clustering analyses on these trajectories depict a clear trend of differential enthalpic affinity of NS3h with ADP, demonstrating a probable mechanism of hydrolysis turnover regulated by the motif-VI loop (MVIL). These findings were experimentally corroborated using viral replicons encoding three mutations at the D471 position. Replication assays using these mutants demonstrated a substantial reduction in viral replication compared to the wild-type. Molecular simulations of the D471 mutants in the apo state indicate a shift in MVIL populations favoring either a closed or open ‘valve’ conformation, affecting ATP entry or stabilization, respectively. Combining our molecular modeling with experimental evidence highlights a conformation-dependent role for MVIL as a ‘valve’ for the ATP-pocket, presenting a promising target for antiviral development.

## Introduction

West Nile Virus (WNV), a member of the *Flavivirus* genus, is a global threat to human health and has become endemic in many areas of the world. ^1,2^ Within the USA, WNV is designated as one of the most important zoonotic diseases.^1,2^ This virus is referred to as a re-emerging pathogenic virus, usually amplified in a mosquito-bird-mosquito enzootic transmission cycle.^3^ However, WNV can infect dead-end hosts, such as humans and horses, causing disease ranging from mild febrile illness to severe neurological disease. Human-human transmission is also reported, for example, through blood transfusion.^4–6^ Currently, there are no FDA-approved vaccines for humans to prevent WNV infection. While a handful of promising human vaccines have been tested, significant barriers exist in advancing these candidates to phase 3 clinical trials. ^7–9^ Thus, the continued development of specific antiviral therapeutics to treat WNV disease is critical. These efforts necessitate a detailed molecular understanding of essential viral protein functions to identify antiviral targets.

The C-terminal end of the non-structural protein 3 (NS3) marks a promising antiviral target due to its pivotal role in viral replication. This region of NS3 functions as a DEAH-box helicase (NS3h) belonging to superfamily-2 (SF2) and is a structurally conserved monomeric protein in the *Flavivirus* genus. This protein is involved in a critical step of replication: unwinding the double-stranded RNA (ds-RNA) replication intermediate by hydrolyzing the nucleotide triphosphate (NTP) in its active site. ^10^ Several studies have reported that increased virulence of WNV strains is due to mutations in the NS3 helicase (NS3h). ^11–14^ WNV NS3h also performs nucleoside 5*^′^*-triphosphatase (NTPase) and 5*^′^*-terminal RNA triphosphatase (RTPase) activities but displays a strong preference for ATP as an NTPase substrate. ^15^ As the binding site of RNA (RNA-cleft) and ATP (ATP-pocket) are within a single monomeric form, it is important to elucidate the structural details of NS3h to understand its function for structure-specific inhibitor design.

The RNA-cleft and ATP-pockets in NS3h are distal and yet functionally correlated. NS3h spans amino acid residues 180 to 619 and adopts a tertiary structure comprised of three domains. Figure 1 provides a graphical illustration elucidating the NS3h structure and its binding sites. Domains I and II are primarily characterized by *β*-strands, sharing the Rossman’s fold, and together they constitute ATP-pocket at the interface. In contrast, domain III is predominantly composed of *α*-helices and forms RNA-cleft along the edges of the other two domains. As in other SF2 helicases, WNV NS3h consists of eight sequence-conserved motifs: motifs **I**, **II**, **III**, and **VI** are involved in ATPase activity, while motifs **Ia**, **IV**, and **IVa** interact with RNA, and motif **V** ^16^ is related to both functions. WNV NS3h can perform ATP hydrolysis and RNA binding independently,^15,17^ but the presence of RNA stimulates ATP hydrolysis, and the unwinding of double-stranded RNA (dsRNA) is an ATP hydrolysis-dependent process.^18,19^ These results indicate that the unwinding process is fueled by the free energy generated through the ATP hydrolysis cycle.

**Figure 1:**
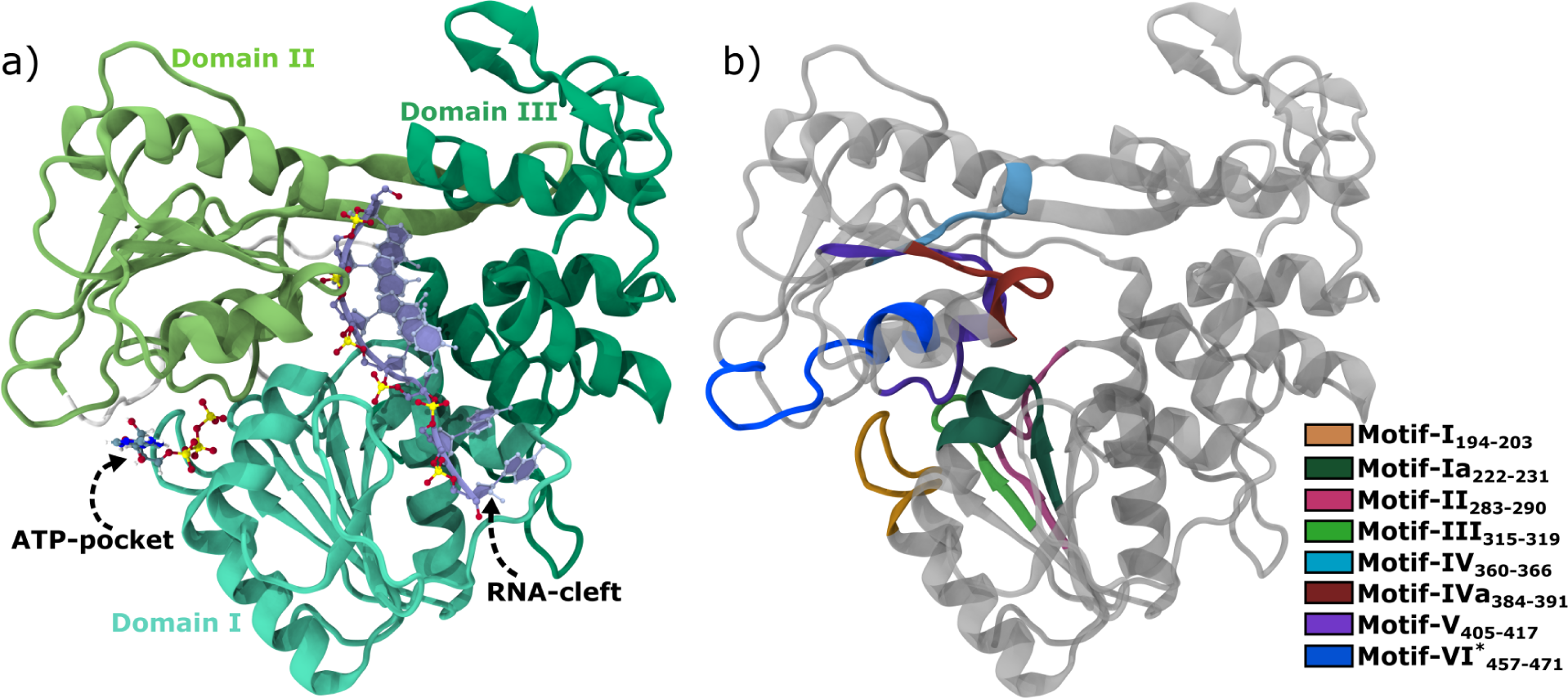
Structural representation of WNV NS3 helicase (NS3h). a) The NS3h structure is displayed in cartoon mode marking three domains with different shade of green color. An ATP-pocket filled with ATP located at the interface of Domain I and Domain II; the RNA-cleft occupied by ssRNA marked at the interface formed by Domain III with Domain I and Domain III. b) Conserved structural motifs are shown here with distinct color code. For clarity, the remaining regions of protein are colored in grey along with removal of ATP and ssRNA. This is an initial modelled structure using DENV4 (PDB ID. 2JLV^20^) and the entire figure prepared by VMD.^21^

Translocation is an elementary step in NS3h-mediated unwinding, and its progression is marked by changing affinity for hydrolysis substrate and product. However, the dsRNA unwinding mechanism of the Flavivirus NS3h remains elusive. A compilation of studies suggest that a small-step, ATP-dependent, recurring translocation cycle leads the protein to move unidirectionally from the 3*^′^* to 5*^′^* end by alternating strong and weak binding to ssRNA.^22–25^ The accumulation of these small steps results in the protein traversing a long stretch of ssRNA, leading to dsRNA unwinding. Experimental studies have observed that ATP binding induces a conformational change in the protein by closing Domain I and Domain II, causing the forward movement of the protein. This suggests that hydrolysis energy does not directly contribute to translocation.^24,26^ Instead, hydrolysis energy leads the protein to stabilize in its new position on ssRNA and return to its initial state for the next cycle. However, the rationale for the effect of hydrolysis energy is based on the conformational change of the protein in the bound product state. Consequently, we propose that hydrolysis alters the affinity of NS3 for products compared to substrates. This shift in affinity within the ATP-pocket creates a differential binding affinity within the RNA-cleft throughout the hydrolysis cycle. Therefore, investigating the NS3h structure while bound to both hydrolysis substrates and products is crucial for identifying the structural changes responsible for this differential nucleotide affinity at the ATP-pocket. These insights could be leveraged to develop strategies to inhibit NS3h function effectively.

Crystal structures of closely related WNV NS3h fail to elucidate the differential affinity of nucleotide (i.e. ADP) throughout the hydrolysis cycle. WNV NS3h shares sequence identity of 69% with its counterparts in DENV and ZIKV. Luo et al.^20^ reported crystal structures of DENV NS3h+ssRNA complex, capturing states of the hydrolysis cycle. These structures reveal notable structural changes in residues coordinating the P*_γ_* and P_i_ moieties, with Lys199 and Glu285 (Lys200 and Glu286 in WNV NS3h respectively) being closer to the P*_γ_* moiety than the P_i_ moiety. In all these structures, P*_β_* moiety of ATP or ADP is coordinated with Arg463 (Arg464 in WNV NS3h). It’s crucial to emphasize that ADP must be released before the subsequent hydrolysis cycle can commence, implying a necessary alteration in ADP affinity—a distinctive feature of motor proteins. However, the crystal structures of DENV alone do not provide insights into this aspect. A similar trend is observed in the crystal structures of ZIKV NS3h when bound to nucleotide and ssRNA, as reported by Yang et al.^27^ and Tian et al.^28^ Nevertheless, a comparison of these crystal structures reveals that ADP shifts outward more significantly in the post-hydrolysis-II state (ADP bound to NS3h+ssRNA) than in the intermediate state, while still interacting with Arg463 (Arg464 in WNV NS3h). Perez-Villa et al. also reported differential affinity for ATP and ADP in their microsecond-long simulations of HCV-NS3h bound to substrates.^29^ This suggests the possibility of another low-affinity state for ADP, which remains unresolved due to the limitations of crystallization techniques for these viral helicases.

Sequence-wise distant to WNV, the tick-borne encephalitis virus (TBEV) NS3h crystal structures underscore the differential affinity based functional mechanism.^30^ As compared to DENV and ZIKV, contrasting structural change is noticed for TBEV NS3h Arg463 (Arg461 for WNV), which switches its direct contact with the P*_β_* moiety in the presence of ATP or ADP+P_i_ to water-mediated contact in the ADP-bound TBEV NS3h. Interestingly, TBEV NS3h Arg466 (Arg464 for WNV) becomes detached from ADP in ADP bound NS3h of TBEV. These observations support the existence of differential nucleotide affinity as a functional mechanism of the NS3 helicase. However, the WNV NS3h shares 49% sequence identity with TBEV, which is lower than that of DENV or ZIKV. Of note, TBEV crystal structures are also unable to depict altered affinity for ADP in the presence of ATP and ADP+P_i_.

We hypothesize that the motif-VI loop plays a significant role in modulating the affinity for nucleotides in hydrolysis states. Arg464 is one of the arginine fingers, with another being Arg461, within **motif-VI** responsible for stabilizing the nucleotide at the ATP-pocket. The structure of **motif-VI** comprises a very short stretch of *α*-helix 11 (*α***11**) and a larger contiguous loop region that connects *α*-helix 11 and *β* strand 14 (*β***14**). To emphasize the loop structure, we have renamed **motif-VI** as **motif-VI loop (MVIL)**. While Arg461 is located in the helix region, Arg464 is situated in the loop region. Biochemical studies report the inhibitory potential of the motif-VI peptide as it is involved in ATP binding, and mutational studies have suggested a conformation-dependent loop function.^31^ In substrate-gated enzymes, the loop structures surrounding the active site have a significant effect, from substrate recognition to catalysis.^32^ Studies demonstrate that loop structures are not merely ornamental connectors of structured regions of proteins; instead, inherent conformational flexibility regulates protein functions, as evidenced in cases such as TPI,^33^ PTPs,^34^ BRD4-1,^35^ etc. In some cases, enzyme activation becomes kinetically slow due to the need to attain the correct conformation of the loop^34^ and strongly dictates the enthalpic contribution to protein-ligand binding affinity.^35^ Taken together, this evidence suggests the need to investigate the nucleotide-specific conformational dynamics of **MVIL**. Here, we report our examination of the differential affinity of NS3h-ADP, focusing on the influence of **MVIL** through theoretical modeling and extensive sampling of substrates bound to WNV NS3h.

## Methods

### Starting Structures

In our study, we modeled substrates bound to WNV NS3h to identify the probable regulatory site of hydrolysis substrate binding and product release along the hydrolysis cycle. However, there are currently no substrates bound WNV NS3h structures available in the Protein Data Bank. WNV NS3h (KUNV) shares 68% sequence identity with DENV4. Therefore, we used DENV4 substrates bound structures (PDB ID: 2JLU, 2JLV, 2JLY, and 2JLZ) as templates to generate ssRNA, ssRNA+ATP, ssRNA+ADP+P_i_, and ssRNA+ADP bound WNV NS3h structures through homology modeling using SWISS-MODEL.^36^ The ssRNA, ATP, ADP+P_i_, ADP, and Mg^2+^ were extracted from the DENV4 structures and manually placed in their respective binding sites on the modeled WNV NS3h structure after alignment with the DENV4 structures. Further, the modeled ssRNA bound WNV NS3h structure was used to prepare the D471E, D471N and D471L structures by changing only the terminal sidechain atoms of respective mutants.

### Simulation Protocol

We conducted all-atom explicit water conventional molecular dynamics (cMD) simulations of the seven modeled structures, i.e., ssRNA, ssRNA+ATP, ssRNA+ADP+P_i_, and ssRNA+ADP-bound WNV NS3h protein along with three mutants (D471E, D471N and D471L) of ssRNA bound NS3h. We utilized the GPU-enabled AMBER18 software^37^ to integrate the equations of motion, employing the ff14SB^38^ and ff99bsc0*χ*OL3^39,40^ force field parameters for the protein and ssRNA, respectively. Similarly, we used ATP,^41^ ADP,^41^ and Mg^2+ 42^ parameters, while P_i_ (H_2_PO_4_)^23^ parameters were adopted from our previous study. We retained the crystallographic water of DENV4 in our modeled structure to maintain the octahedral organization of Mg^2+^ surrounded by ‘O’ atoms. The neutral pH protonation state was assigned to amino acids based on the PropKa^43^ generated pKa values. Special attention was given to histidine residues for assigning protonation states, specifically HIE194, HID251, HID262, HIE267, HID273, HID287, HIE477, HIE487, HIE558, and HIE604. The protein complexes were solvated with TIP3P^44^ water model in a cubic box, leaving a 20 Å buffer length between the protein and the box edge. Sodium (Na^+1^) and Chloride (Cl^−1^) ions were added to neutralize the charge and maintain an ionic strength of 0.15M in the solution using the ‘addIons2’ module of tleap. The resulting box volume was 107 Å^3^.

Each system was subsequently minimized in seven steps by gradually decreasing the restraint on the protein and substrates (including Mg^2+^) to maintain initial coordination, with each step running for 2000 steps of steepest descent. The restraint force constants used were 150, 100, 50, 10, 1, and 0.1 kcal·mol^−1^ · Å^−2^ for steps one through six, respectively. Subsequently, we gradually heated the system from 0 K to 300 K over 500 ps in an NVT ensemble by applying a restraint of 150 kcal·mol^−1^ · Å^−2^ on the protein complex, followed by pressure (1 bar) equilibration for 1.4 ns in seven steps, gradually reducing the restraint force to relax the system. Finally, we performed a production run for 5 *µ*s and 3 *µ*s for wild-types and mutants respectively and generated conformations in an NPT ensemble. Periodic boundary conditions (PBC) were applied in all directions. The Langevin dynamics thermostat and Monte Carlo barostat were employed to maintain the systems at 300 K and 1 bar. Direct nonbonding interactions were calculated up to a 12 Å distance cutoff. The SHAKE algorithm was used to constrain covalent bonds involving hydrogen.^45^ The particle-mesh Ewald method was utilized to account for long-ranged electrostatic interactions.^46^ A 2 fs integration time step was applied, with energies and positions recorded every 2 ps. In total, we conducted approximately 20 *µ*s of production simulations.

### Cell Lines

293T cells were acquired from ATCC and propagated in DMEM supplemented with 10% heat-inactivated FBS and 1% GlutaGro (Gibco). Cell lines were validated to be free of Mycoplasma contamination using a Mycostrip kit (Invitrogen).

### Plasmids and Generation of Mutants

The lineage II WNV Replicon encoding GFP was a generous gift from Dr. Ted Pierson at the NIH.^47^ To generate mutants, primers were ordered from Integrated DNA Technologies (IDT) encoding D *→* E, D *→* N or D *→* L mutations at position 471 in NS3. Site-directed mutagenesis was performed using InFusion cloning (Takara). The sequence of all replicon plasmids was validated using Oxford nanopore sequencing (Plasmidsaurus).

### Cell Transfection and Replicon Assays

293T cells were seeded into 6-well plates and transfected with replicon plasmid using Xtreme-Gene9 transfection reagent (Sigma). 24 hours post-transfection, cells were replated in duplicate 6-well plates for RNA and protein analysis and 96-well plates for automated immunofluorescence analysis. 48 hours post-transfection cells were collected in Tri-reagent (Zymo) or RIPA buffer supplemented with protease inhibitors and 96-well plates were fixed with 4% paraformaldehyde.

### cDNA generation and qRT-PCR

RNA was extracted from Tri-reagent using the Direct-Zol RNA extraction kit (Zymo). cDNA was generated using total RNA and M-MLV reverse transcriptase (Invitrogen). qPCR was performed to measure 18s rRNA and WNV replicon RNA against a standard curve of a pooled reference to calculate relative RNA abundance.

### Western blotting

Cell lysates were freeze-thawed to ensure complete lysis, then spun at 21,000xg for 15*^′^* at 4*^◦^*C to remove cell debris. Lysates were transferred to fresh tubes, mixed with 6x Laemmli buffer, and heated at 95*^◦^*C for 5*^′^*. 20mg protein lysates were resolved on 10% tris/glycine polyacrylamide gels and transferred to PVDF (0.45 *µ*m pore) before being blocked in 5% milk in TBST. Blots were incubated in a-Actin, a-GFP, and a-WNV NS3 antibodies overnight, then washed with TBST and incubated with HRP-conjugated secondary antibodies, washed, and developed using ECL (Cytiva).

### Immunofluorescence

After paraformaldehyde fixation, plates were washed 3x in PBS and incubated with Hoechst (5mg/mL) for 1 hour at RT. Plates were washed 3x in PBS and imaged using an ImageXpress Pico automated imaging system to quantify total cells and GFP expression.

### Statistical Analysis

Statistical analyses were performed using GraphPad Prism 10. One-way ANOVA were performed with Dunnett’s correction for multiple comparisons.

### Primers for Cloning and RT-qPCR

<u>D471E-F</u> AGTTGGTGAGGAGTATTGCTATGGAGGGCACAC

<u>D471E-R</u> TACTCCTCACCAACTTGTGATGGGTTTCTTCC

<u>D471N-F</u> AGTTGGTAACGAGTATTGCTATGGAGGGCACAC

<u>D471N-R</u> TACTCGTTACCAACTTGTGATGGGTTTCTTCC

<u>D471L-F</u> AGTTGGTCTGGAGTATTGCTATGGAGGGCACAC

<u>D471L-R</u> TACTCCAGACCAACTTGTGATGGGTTTCTTCC

<u>18s-F</u> GGCCCTGTAATTGGAATGAGTC

<u>18s-R</u> CCAAGATCCAACTACGAGCTT

<u>WNVII NS1-F</u> GGAGGTAGAGGACTTTGGATTTG

<u>WNVII NS1-R</u> GCACAGCCATGTTGTTCTTG

### Antibodies

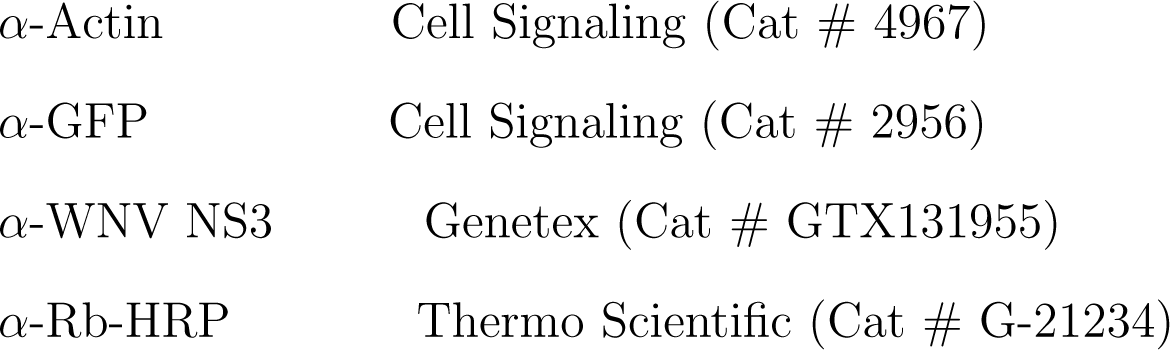

### Analyses

#### Clustering

In this study, we employed the Size-and-Shape Space Gaussian Mixture Model (Shape-GMM)^48^ to identify structural states (or clusters) of the **MVIL**. This model determines optimal parameters for multivariate Gaussians using particle positions as features. To determine the appropriate number of clusters, we utilized the elbow heuristic coupled with cross-validation (CV). The elbow heuristic suggests choosing a number of clusters at which point the log-likelihood as a function of number of clusters does not increase significantly. This is best quantified by the minimum in the second derivative of log-likelihood as a function of numebr of clusters. CV is used to ensure that the model has not been overfit. Five training sets for all systems were determine sampling error. The assignment of a frame to a cluster was achieved by minimizing the Mahalanobis distance after a weighted alignment. The kronecker product model of the covariances was used in all shapeGMM analyses.

#### Linear Discriminant Analysis (LDA)

LDA is a dimensionality reduction algorithm for cluster separation that works by minimizing intra-cluster variance while maximizing inter-cluster variances. LDA on shapeGMM derived clusters has been found to produce good estimates of reaction coordinates separating the clusters.^49^ This supervised technique utilizes globally aligned and cluster-labeled particle positions to learn the cluster-separating vectors. LDA generates K-1 vectors that effectively best separate the clusters where K denotes number of clusters identified from shape-GMM. We employed the Python library scikit-learn with the single-value decomposition (SVD) method to handle variance.

#### Binding Enthalpy

In our NPT simulation, the binding enthalpy of the system is determined as the interaction energy between substrate (i.e., ADP) and protein. We consider the non-bonded interactions, including electrostatic and van der Waals interactions. We used Cpptraj *lie* module to calculate the binding enthalpy with a long-range cutoff of 12 Å.

### Model Corroboration

Closest and conserved contact analysis of Mg^2+^, hydrolysis substrates, and ssRNA demonstrates well-modeled NS3h bound substrates and stable simulations. The Mg^2+^ ions form octahedral coordination with oxygen atoms. We computed Mg^2+^-coordinating oxygen atoms within 2.5 Å, and the results are tabulated in Table S1. We observed that the octahedral arrangement is maintained throughout the simulation for all hydrolysis states. The Mg^2+^-T201 coordination is present in all hydrolysis states, while E286 is only part of the coordination shell in the ATP bound state. The participation of ATP or ADP oxygens in the coordination shell of Mg^2+^ decreases from the ATP-bound state to the ADP-bound state. Similarly, the Mg^2+^-surrounding water molecules increase from the ATP-bound to the ADP-bound NS3h. We also calculated the occurrence probability of conserved residues in the ATP-pocket (Table S2) and RNA-cleft (Table S3) that form contacts with ATP/ADP+P_i_/ADP and ssRNA, respectively (see the table). Most of the contacts show a 99% existence probability, with only a very few contacts at 70%. However, the probability of contacts varies significantly between the hydrolysis states.

## Results and Discussion

We present and discuss our in-depth analysis of the differential enthalpic affinity of WNV NS3 helicase for nucleotide hydrolysis, a critical factor in the hydrolysis turnover process that is essential for motor protein function. ATP hydrolysis involves a multistage process characterized by the changing identity of the nucleotide at the ATP-pocket. Based on available crystal structures of substrates bound to NS3h, we can divide the hydrolysis cycle into four stable states.^20,27,28,30^ For the sake of clarity, we will refer to these states as hydrolysis states. These states include the apo state, where the ATP-pocket is vacant and only ssRNA is present at the RNA-Cleft of NS3h (ssRNA); the pre-hydrolysis state, where ATP enters the ATP-pocket of NS3h, transitioning it into the catalytic active form (ssRNA+ATP); the post-hydrolysis-I state, where NS3h is bound to the products (ADP and P_i_) of ATP hydrolysis (ssRNA+ADP+P_i_); and finally, the post-hydrolysis-II state, where NS3h is bound with ADP (ssRNA+ADP). After ADP release, NS3h returns to the apo state in preparation for the next cycle. Since ADP is a common nucleotide among the hydrolysis states, we present our analysis of nucleotide-protein affinity focusing on ADP (in case of ATP, we do not consider P*_γ_* moiety). For clarity, we organize our results into four sections. In the first section, we report the predominant role of **MVIL** in altering the affinity of NS3h-ADP for hydrolysis states. The second section describes the conformational plasticity of **MVIL** and nucleotide-dependent sampling of **MVIL**. In the third section, we delve into a more detailed investigation of substrate-specific **MVIL** conformations and their regulatory role in nucleotide entry and exit. The last section describes results of targeting D471 of **MVIL**, which indirectly and significantly affected viral replication.

### Motif-VI Loop Plays a Predominant Role in ATP-Dependent Differential Binding Enthalpy of NS3h-ADP

Experimental reports have been unable to elucidate the differential affinity of ADP in each hydrolysis state. Structural, biochemical, and mutagenesis studies suggest that motifs **I**, **II**, **III**, and **VI** are involved in the ATPase activity of NS3h.^10^ Crystallographic structures have reported ssRNA induced structural changes in motif **I** and alterations in coordination, mostly in the P*_γ_* moiety.^20^ However, limited attention has been given to the changes in ADP affinity. The crystallographic structures of ZIKV and DENV have failed to provide differential coordination for ADP in various hydrolysis states.^20,28^ While the structures of TBEV shed light on motif **VI** mediated ADP coordination changes in the post-hydrolysis-II state, they also fall short in explaining affinity changes in the post-hydrolysis-I state compared to the pre-hydrolysis state.^30^ These crystallographic reports are based on single structures bound to nucleotide and lack insights from conformational ensembles.

WNV NS3h alters its enthalpic affinity for ADP according to the hydrolysis state as a functional mechanism of ATP-hydrolysis turnover. In Figure 2(a), we present the distribution of interaction energy of the entire NS3h protein and ADP (E^NS3h–ADP^) for ss-RNA+ATP, ssRNA+ADP+P_i_, and ssRNA+ADP systems. This interaction energy can approximate the binding enthalpy, disregarding the pressure-volume terms and change in internal energy of NS3h and ADP. These distributions are statistically distinct, as quantified by a t-test (Table S4). Compared to the pre-hydrolysis state, the binding enthalpy distribution of the post-hydrolysis-I state is left-shifted, indicating a stronger affinity. The post-hydrolysis-II state binding enthalpy is right-shifted compared to other states, indicating a weaker affinity of NS3h for ADP. In Figure 2(a), the mean and standard error are also annotated for each distribution.

**Figure 2:**
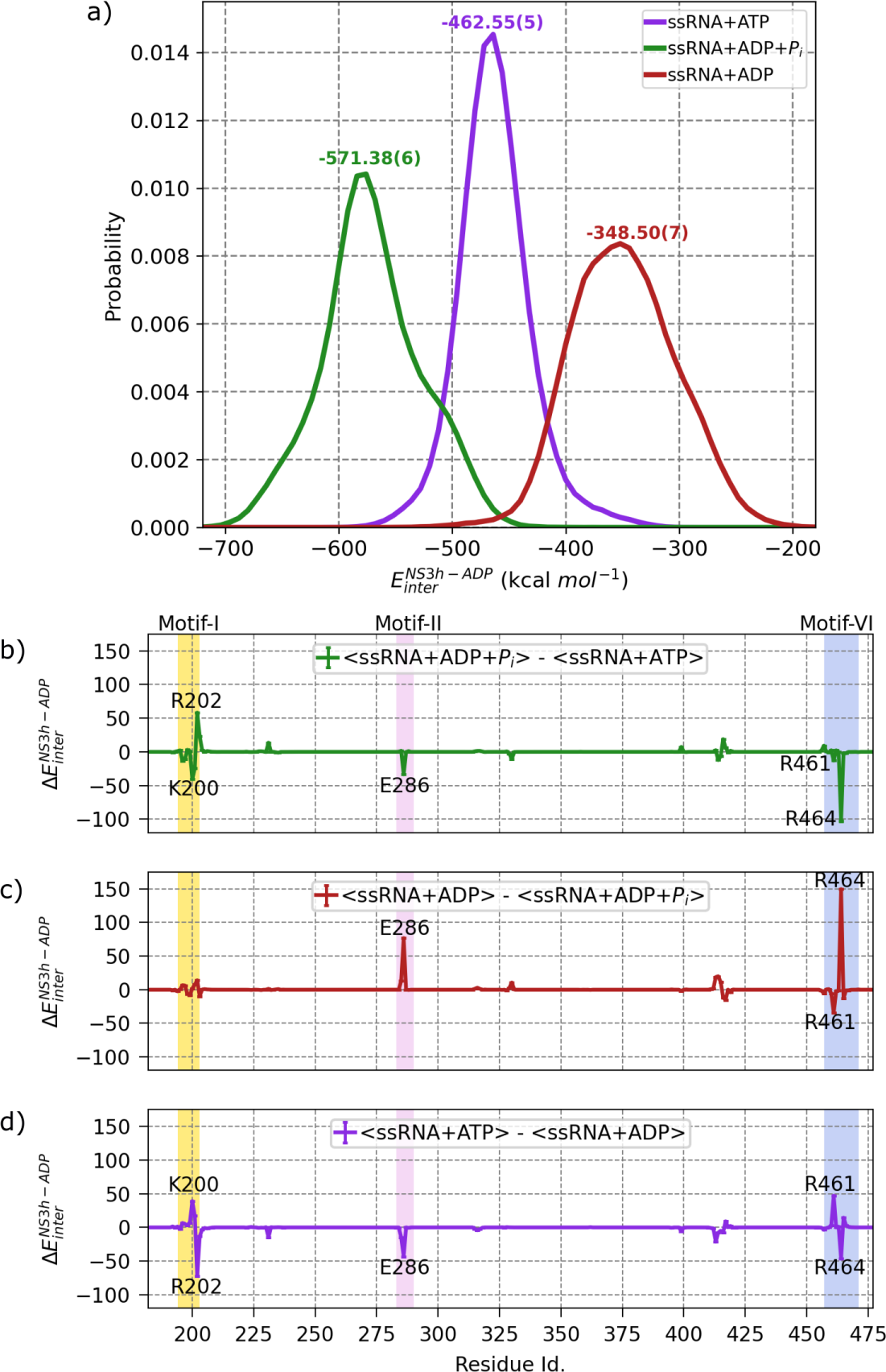
Nucleotide-dependent alteration of binding enthalpy of WNV NS3h and ADP along the hydrolysis cycle. a) The binding enthalpy correspond to non-bonded linear interaction energy (E_inter_) of NS3h and ADP and here, presented as a probability distribution for ssRNA+ATP (pre-hydrolysis), ssRNA+ADP+P_i_ (post-hydrolysis-I) and ss-RNA+ADP (post-hydrolysis-II) states. For the ssRNA+ATP system, we excluded the P*_γ_* moiety in the calculation. The interaction energy calculation is performed over the entire 5*µ*s of production run for each simulated system with *lie* module of *cpptraj*. Further, normalized histogram is calculated for 100 bins with same lower and upper bound in each system using python (*numpy.histogram*). In the figure, the mean value of each distribution is annotated by respective color with standard error of mean for last digit in parentheses. The difference of NS3h-ADP binding enthalpy between the states decomposed into protein residues which are presented as differential binding enthalpy for b) ssRNA+ADP+P_i_ (post-hydrolysis-I) and ssRNA+ATP (pre-hydrolysis), c) ssRNA+ADP (post-hydrolysis-II) and ssRNA+ADP+P_i_ (post-hydrolysis-I), d) ssRNA+ATP (pre-hydrolysis) and ssRNA+ADP (post-hydrolysis-II). The ΔE_inter_ of a residue is computed by subtracting the mean of interaction energy of residue and ADP of corresponding substrates bound helicase. The error is propagated by following addition rule and presented as error-bar. *Matplotlib* is used to generate the plot.

The highest affinity in the post-hydrolysis-I state suggests that the helicase is enthalpically favored for ATP hydrolysis. Furthermore, the hydrolysis of the ATP substrate induces conformational changes in the protein, strengthening its grip on ADP. The strongest affinity for ADP in the post-hydrolysis-I also suggests that ADP is unable to release in this state, unless there is the presence of another dominating force. Transitioning to the post-hydrolysis-II state, the weakest affinity indicates that the protein has a weaker hold on ADP, facilitating its release. The enthalpic energy difference of 222.88 kcal mol^−1^ between the post-hydrolysis-II and post-hydrolysis-I state suggests that the presence of P_i_ impedes the protein’s transition from the post-hydrolysis-I to the post-hydrolysis-II state enthalpically. We hypothesize that P_i_ release may facilitate this transition, a process often recognized as a rate-limiting step in motor proteins, as suggested in the literature.^50–52^ ADP release resets the protein to its initial apo state, allowing NS3h to bind ADP more strongly in the pre-hydrolysis state for the subsequent cycle than in the post-hydrolysis-II state of the previous cycle. It is important to note that the stronger affinity of NS3h for ADP in the pre-hydrolysis and post-hydrolysis-I states also reflects the influence of P*_γ_* and P_i_, respectively. Nevertheless, the calculation of NS3h and ADP binding enthalpy has facilitated the comparison of these intriguing insights across all hydrolysis states.

Motif-VI loop (MVIL) emerges as a central player orchestrating the modulation of binding enthalpy between NS3h and ADP throughout the hydrolysis cycle. As illustrated in Figure 2(b-d), we delve into the affinity of each protein residue for ADP, examining the differential binding enthalpy. Progressing through the hydrolysis cycle, we explore three transitions: the upper panel represents changes from the pre-hydrolysis state to the post-hydrolysis-I state, the middle panel delves into shifts from the post-hydrolysis-I state to the post-hydrolysis-II state, and the lower panel examines the transitions from the previous post-hydrolysis-II state to the next pre-hydrolysis state. In this analysis, K200, R202, E286, R461, and R464 exhibit significant alterations in their affinity for ADP among the hydrolysis states.

K200 and R202, situated within **motif-I** (P-loop), exhibit interesting motions. When the protein shifts from the pre-hydrolysis state to the post-hydrolysis-I state (Figure 2(b)), K200 fosters an attractive affinity, while R202 exerts a repulsive influence on ADP. Conversely, during the transition from the post-hydrolysis-II state (previous hydrolysis cycle) to the pre-hydrolysis state of next cycle (Figure 2(d)), we observe a complete reversal in trends, both in direction and magnitude. This suggests that K200 plays a crucial role in hydrolysis, while R202 contributes to stabilizing ADP (or ATP) in the P-loop during the pre-hydrolysis state. However, intriguingly, no significant changes emerge in the post-hydrolysis-II state compared to the post-hydrolysis-I state (Figure 2(c)), implying that K200 and R202 play a lesser role during this phase.

E286, a crucial catalytic residue, displays varying ADP affinities in hydrolysis states. It exhibits attraction when transitioning from pre-hydrolysis to post-hydrolysis-I (Figure 2(b)), switches to repulsion in post-hydrolysis-II compared to post-hydrolysis-I (Figure 2(c)), and turns attractive again when moving from post-hydrolysis-II to pre-hydrolysis (Figure 2(d)). This shift is due to the position of E285 relative to ADP; it is distant when hydrolysis is required, creating attraction. In the absence of P_i_, E286 moves closer in the post-hydrolysis state-II, leading to repulsion.

Significantly, the trend of hydrolysis state-dependent changes in binding enthalpy highlights the prominent roles of **MVIL** residues, R461 and R464, in altering the overall binding enthalpy. As the protein moves from pre-hydrolysis to post-hydrolysis-I state (Figure 2(b)), R464 strongly binds to ADP, resisting its release, while R461 shows minimal change due to its close coordination with P*_γ_* or P_i_. Transitioning to the post-hydrolysis-II state, R464 exhibits a significant change, shifting towards repulsion (Figure 2(c)). This change is likely driven by the absence or release of P_i_, preparing the protein for recycling by releasing ADP. In the absence of P_i_, R461 moves closer to ADP, creating an attractive interaction. During the next cycle, R461 becomes repulsive, while R464 becomes attractive in the pre-hydrolysis state compared to the post-hydrolysis-II state (Figure 2(d)), stabilizing ADP (part of ATP) for access of P*_γ_* for catalysis.

R461 and R464 in the **MVIL** exhibit localized correlations. In Figure S1 we plotted the correlation of R461 (a) and R464 (b) with other residues of the protein. Notably, R464 exhibits a remarkably strong correlation, particularly in the absence of nucleotide at the ATP-binding pocket (apo state) and to smaller extent in the post-hydrolysis states. Conversely, the pre-hydrolysis state shows the least correlation. These correlations signify the conformational adaptability of the **MVIL** in the presence of nucleotide, which may play a crucial role in modulating its affinity for ADP.

### Conformational Induction of Motif VI Loop Along the Hydrolysis Cycle

The conformational dynamics of active site loops play a crucial role in catalysis, impacting substrate-specific interactions and loop-mediated processes. These loops, previously viewed as structural elements, are now recognized as functional contributors, capable of various roles beyond mere structural connections. Their inherent flexibility allows them to perform diverse functions and, often, to act as allosteric modulators. ^53^ Notably, the active site loop’s conformational variations^34,54^ influence substrate selection, intermediate state stabilization, and product release. In some enzymes, the sampling of catalytically active conformations is a rate-limiting step due to the wide range of possible conformations, affecting the catalytic rate. This conformational flexibility also leads to sampling of catalytically inactive states, potential targets for disrupting function through subtle changes.^34,55,56^ Additionally, substrate presence can tune loop conformations throughout the catalysis cycle, transforming a flexible loop into a rigid one.^57^ This substrate-dependent conformational change can be explained thermodynamically by either a *selection* or *induction* mechanism, involving the sampling of existing protein conformations or unique conformations, respectively. ^58^ Distinguishing between these mechanisms is challenging, as both can occur depending on substrate concentration. A deeper understanding of these processes holds promise for targeted interventions in ATP hydrolysis.

Identifying unique conformations of the MVIL loop within the ensemble is crucial for understanding its regulatory role in the differential affinity for ADP throughout the hydrolysis cycle. This loop surrounds the active site (ATP-pocket) and is located opposite to the P-loop. Evolutionary, loop sequences are highly conserved,^32^ particularly in the WNV motif VI loop (residues 461 to 472), which shows significant conservation among *Flavivirus* members. The strict conservation of the MVIL sequence underscores its importance in ATP hydrolysis. Therefore, it is vital to categorize the explored conformational ensemble of the motif VI loop and identify conformations specific to each hydrolysis state. This knowledge can be valuable in understanding how conformational dynamics alter the affinity of NS3h for ADP.

We identified five unique clusters (or states) within the conformational ensemble of the motif VI loop using the weighted Shape-GMM (W-SGMM) clustering algorithm. In Figure 3(a), we plotted the log likelihood per frame as a function of the number of clusters during a scan. This calculation was performed on the combined trajectories from apo (ssRNA), pre-hydrolysis (ssRNA+ATP), post-hydrolysis-I (ssRNA+ADP+P_i_), and post-hydrolysis-II (ssRNA+ADP) state simulations, totaling approximately 2 × 10^3^ frames. We focused on the heavy atoms of the **MVIL**. For the scan, we selected five training sets, each consisting of randomly chosen 1 × 10^3^ frames. The W-SGMM was fitted to the training set (blue curve), and cross-validation was performed on the remaining 1 × 10^3^ frames (orange curve). Both the W-SGMM training and cross-validation (green curve) exhibited a significant change in slope at five clusters, indicating the presence of unique clusters. After confirming the presence of five unique clusters, we applied this clustering to the entire combined trajectory, and representative structural images for each cluster are shown in Figure 3(b). To further explain the uniqueness of these clusters, we projected the MVIL structures onto a 2D free energy surface using LD1-LD2 (Figure 4) and LD3-LD4 (Figure 5). The LD1 vector distinctly separated C1, C2, and C4 clusters, while LD2 separated C1 and C2. LD3-LD4 separation encompassed C1, C2, C3, and C5 clusters.

**Figure 3:**
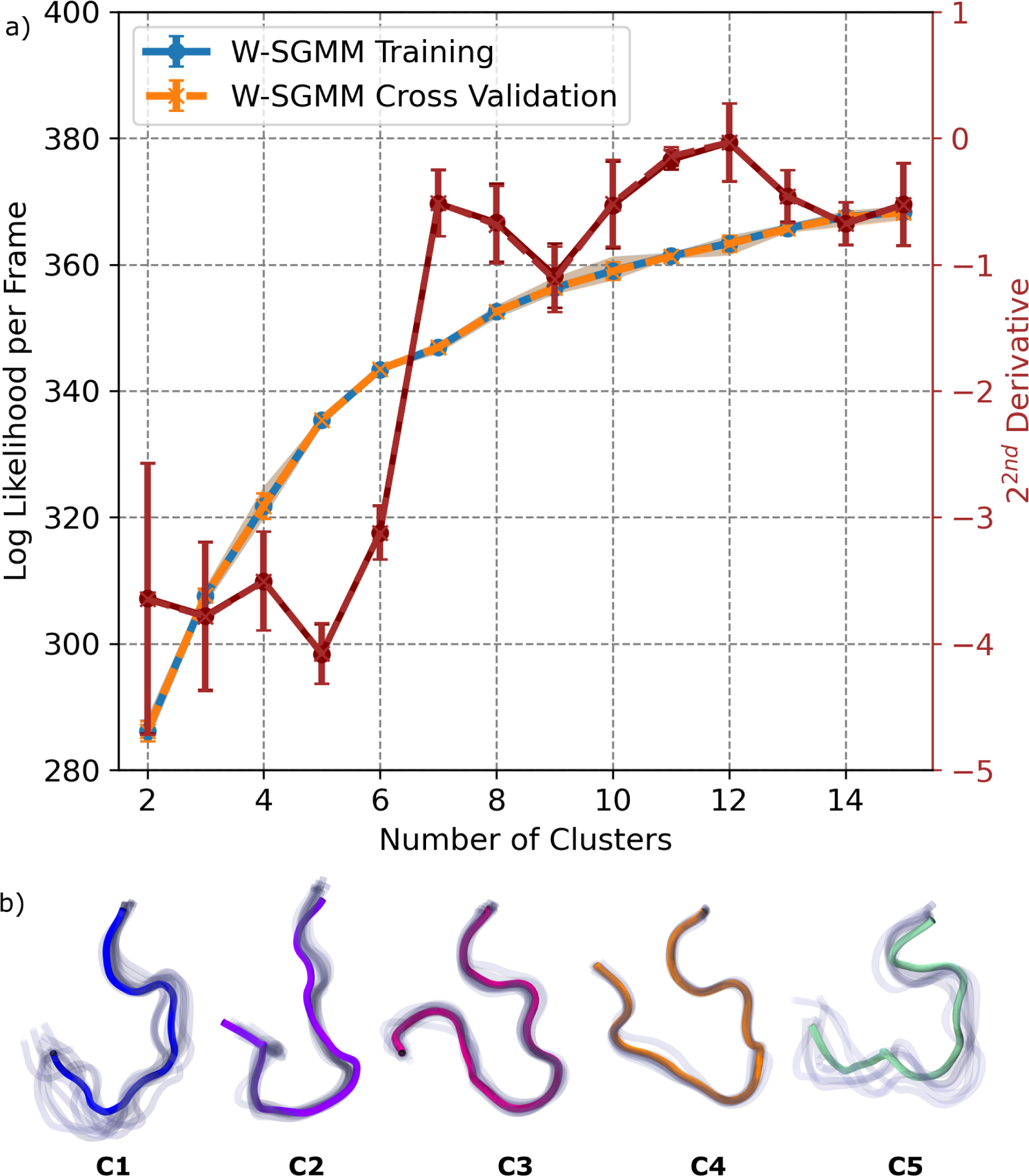
Clustering of motif-VI loop ensembles sampled in the hydrolysis states of WNV NS3h. a) The log likelihood per frame as a function of clusters is plotted here with change of slope measure by 2^nd^ derivative. The clustering is performed with Shape-GMM algorithm by combining trajectories of simulated systems here. In our calculation, the feature consists with positions of all heavy atoms of residues 461 to 472. In this calculation, randomly 1 × 10^3^ frames are selected for training while prediction done on 1 × 10^3^ frames; each training run is iterated for 5 times for each cluster. For change in slope, 2^nd^ derivative is averaged over all iterations of CV scan and accordingly error is calculated. b) Representation of five motif-VI loop clusters based on the minima of slope change of log liklihood per frame. In each cluster, ten (10) randomly selected frames are displayed in translucent while the average structure is highlighted.

**Figure 4:**
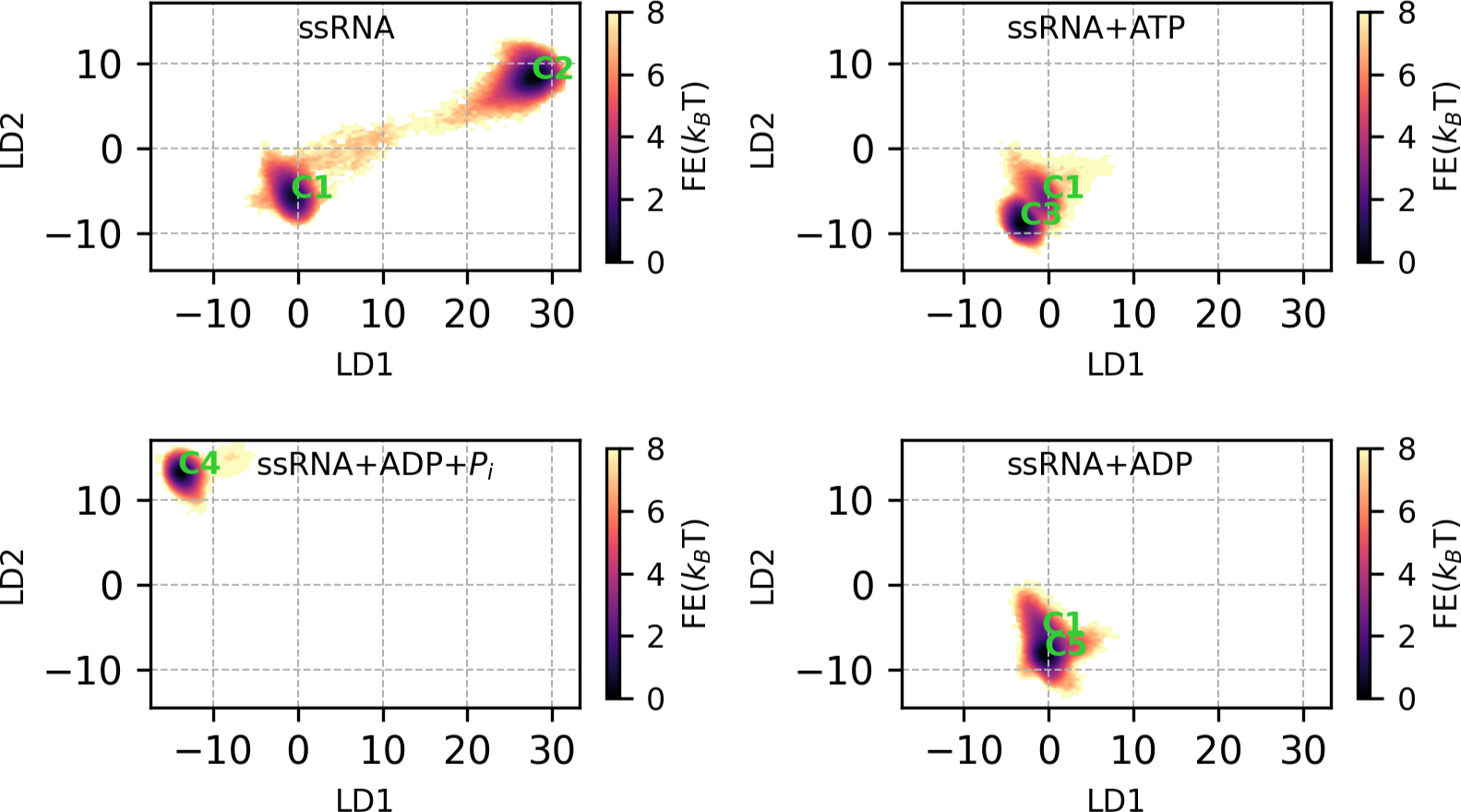
Free energy surface of motif-VI loop clusters in LD1 and LD2 space along the hydrolysis cycle. The free energy surface is presented over the LD1 and LD2 vectors computed by linear discriminant analysis (LDA) on the clustering identity tagged motif-VI loop conformational ensembles. Later, conformations belonging to ssRNA, ssRNA+ATP, ssRNA+ADP+P_i_ and ssRNA+ADP are projected along the LD1 and LD2 vectors followed by 2D histogram.

**Figure 5:**
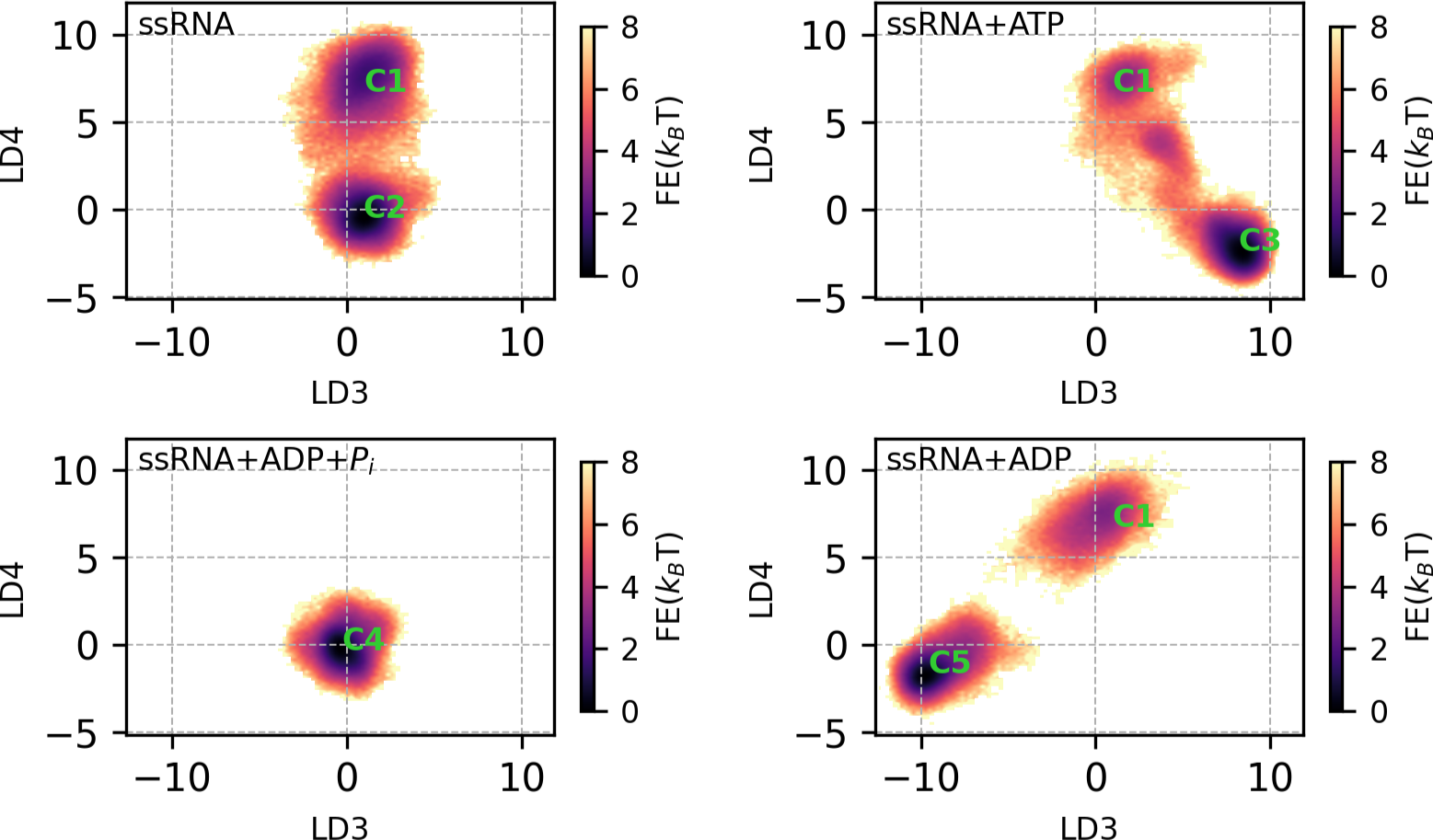
Free energy surface of motif-VI loop clusters in LD3 and LD4 space along the hydrolysis cycle. The free energy surface is presented over the LD3 and LD4 vectors computed by linear discriminant analysis (LDA) on the clustering identity tagged motif-VI loop conformational ensembles. Later, conformations belonging to ssRNA, ssRNA+ATP, ssRNA+ADP+P_i_ and ssRNA+ADP are projected along the LD3 and LD4 vectors followed by 2D histogram.

**Table 1:** ATP-dependent Motif-VI loop cluster probability. These clusters are identified by shape-GMM clustering. Along the hydrolysis cycle, the presence or absence of hydrolysis substrate and products induced MVIL to predominately sample unique cluster.

| Clustrs | ssRNA | ssRNA+ATP | ssRNA+ADP+P<sub>i</sub> | ssRNA+ADP |
| --- | --- | --- | --- | --- |
| **C1** | 0.33 | 0.12 | – | 0.16 |
| **C2** | **0.67** | – | – | – |
| **C3** | – | **0.88** | – | – |
| **C4** | – | – | **1.0** | – |
| **C5** | – | – | – | **0.84** |

Nucleotide-dependent **MVIL** clusters suggest conformational induction at play during the hydrolysis progression. In Table 2, we present cluster probabilities for the apo (ssRNA), pre-hydrolysis (ssRNA+ATP), post-hydrolysis-I (ssRNA+ADP+P_i_), and post-hydrolysis-II (ssRNA+ADP) states. In the apo state, C1 and C2 clusters are sampled, with C2 being more prevalent. Upon ATP binding in pre-hydrolysis state, the sampling of C1 is reduced, but C2 is not sampled. Instead, the pre-hydrolysis state predominantly samples the C3 cluster of the **MVIL**. After hydrolysis, in the post-hydrolysis-I state, we do not observe the presence of C1, C2, or C3 clusters but only sample the C4 cluster. In the post-hydrolysis-II state, we notice the reappearance of the C1 cluster with smaller probability, but C5 is the most frequently sampled cluster.

**Table 2:** ATP-dependent R461(NH1) formed contacts distance and its deviation within the clusters. Presented values denote average of the cluster in the respective hydrolysis state while parentheses value denote standard error of mean for last digit. Units is Å.

| Contact | ssRNA |  | ssRNA+ATP |  | ssRNA+ADP+P<sub>i</sub> | ssRNA+ADP |  |
| --- | --- | --- | --- | --- | --- | --- | --- |
|  | C1 | C2 | C1 | C3 | C4 | C1 | C5 |
| E286(OE1) | 9.478(3) | 4.769(2) | 8.282(4) | 6.176(2) | 10.524(2) | 2.936(2) | 3.050(1) |
| E413(OE1) | 10.971(1) | 9.106(1) | 10.128(3) | 10.416(1) | 10.234(0) | 9.703(3) | 6.504(3) |
| N417(OD1) | 6.920(2) | 4.207(2) | 4.706(5) | 6.233(2) | 5.374(2) | 8.404(4) | 10.275(2) |
| Q457(OE1) | 8.047(3) | 4.780(1) | 5.466(4) | 6.125(1) | 5.288(0) | 4.730(1) | 5.871(2) |
| P<sub>γ</sub> | – | – | 3.946(2) | 3.377(0) | – | – | – |
| P<sub>i</sub> | – | – | – | – | 7.249(1) | – | – |
| P<sub>β</sub> | – | – | 6.316(2) | 6.050(0) | 6.186(0) | 5.197(2) | 4.594(1) |

It is important to note that each hydrolysis state displays the presence of a unique **MVIL** cluster. Our understanding of this differential sampling in line with the hydrolysis cycle is as follows: in the pre-hydrolysis state, the C1 cluster is selected for stabilizing the ATP binding at the ATP-pocket, and the binding induces C2 to provide a strong grip on ADP. We hypothesize that C2 represents the catalytic active state. Transitioning to the post-hydrolysis-I state, the presence of products induces the sampling of C4, which forms the strongest grip on ADP, preventing its release. After the release of P_i_, it induces the sampling of C5 in the post-hydrolysis-II state to weaken the grip on ADP, along with selecting the C1 cluster. This suggests that the preparation for the next hydrolysis cycle begins before the end of the previous cycle. ADP release induces C2 sampling, and in the absence of any nucleotide at the ATP-pocket, it leads to the sampling of C1 to prepare for the next hydrolysis cycle.

### Motif-VI Loop Acts as a Valve for the Nucleotide

In this section, we delve into the molecular mechanisms underlying NS3h-ADP affinity, particularly focusing on the key residues R461 and R464 within the **MVIL**. Our approach involves categorizing protein conformations based on **MVIL** cluster identity, hydrolysis state dependent separation, aligning them with the respective cluster means and co-variances of **MVIL**, and assessing the distances to the closest neighboring residues of R461 and R464 within a 5 Å radius. For clarity, we present our findings separately for each hydrolysis state and provide visual representations of MVIL clusters and their neighboring residues in Figure 6. Detailed distance values for R461 and R464 are available in Tables 2 and 3, respectively, facilitating a comprehensive understanding of the molecular interactions involved.

**Figure 6:**
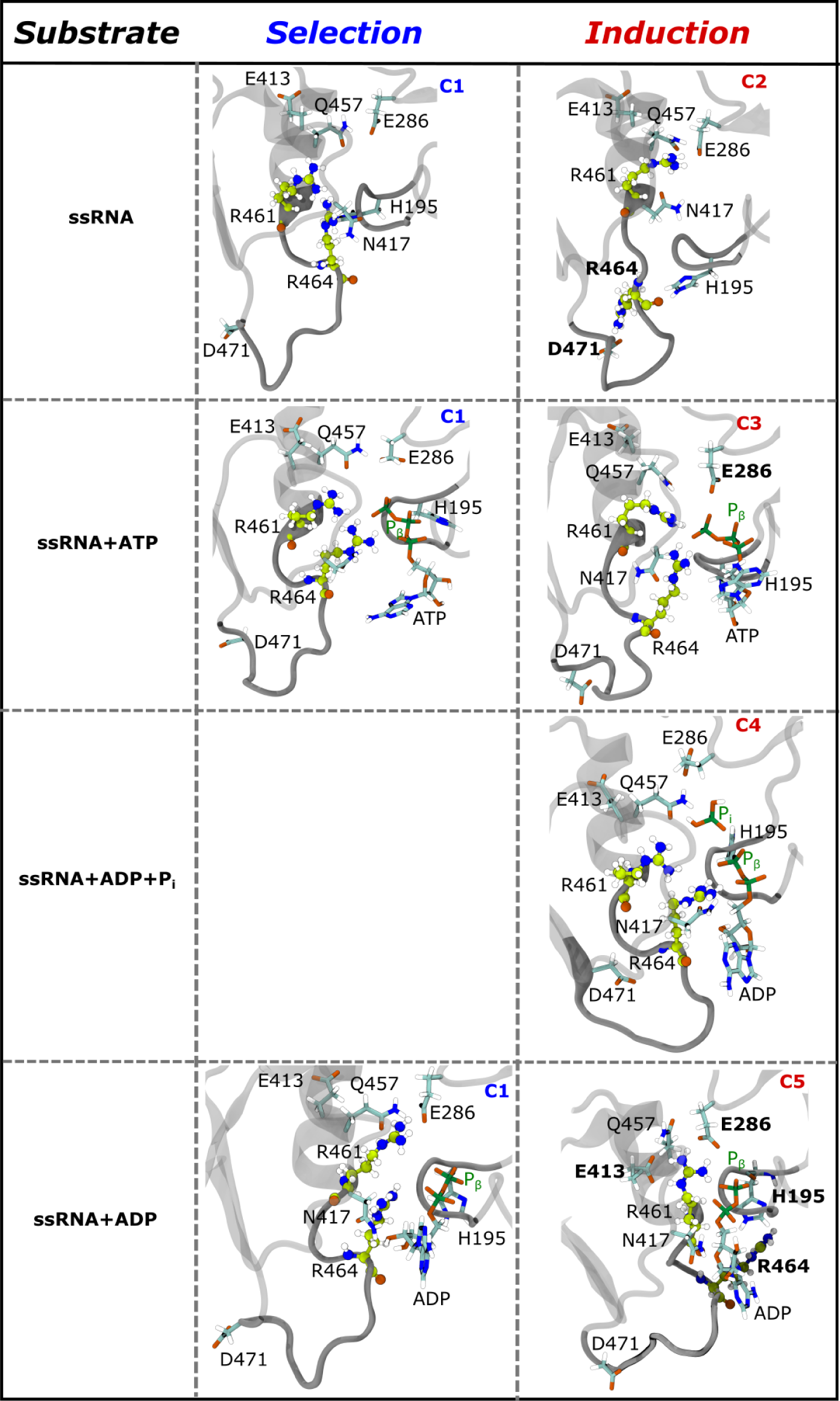
Nucleotide-dependent conformational *selection* or *induction* mechanism of MVIL. Here, we are understanding the role of **MVIL** clusters by evaluating R461 and R464 motion. Based on the clustering identity of **MVIL**, entire substrates bound protein conformations are extracted. R461 and R464 colored in green while closest contacts of these residues are presented in cyan. From row 1 to row 4, nucleotide dependent **MVIL** conformations are presented for ssRNA (apo), ssRNA+ATP (pre-hydrolysis), ssRNA+ADP+P_i_ (post-hydrolyis-I) and ssRNA+ADP (post-hydrolyis-II) states respectively

**Table 3:** ATP-dependent R464(NH2) formed contacts distance and its deviation within the clusters. Presented values denote average of the cluster in the respective hydrolysis state while parentheses value denote standard error of mean for last digit. Units is Å.

| Contact | ssRNA |  | ssRNA+ATP |  | ssRNA+ADP+P<sub>i</sub> | ssRNA+ADP |  |
| --- | --- | --- | --- | --- | --- | --- | --- |
|  | C1 | C2 | C1 | C3 | C4 | C1 | C5 |
| H195(ND1) | 7.096(4) | 13.024(3) | 10.617(4) | 10.466(2) | 11.553(2) | 5.954(6) | 4.479(2) |
| N417(OD1) | 7.026(4) | 17.933(3) | 5.229(6) | 4.530(1) | 5.091(2) | 11.270(8) | 14.734(4) |
| D471(OD1) | 20.538(4) | 4.240(3) | 17.655(8) | 18.977(2) | 11.161(1) | 16.606(8) | 20.073(3) |
| P<sub>γ</sub> | – | – | 3.784(1) | 3.797(0) | – | – | – |
| P<sub>i</sub> | – | – | – | – | 6.673(0) | – | – |
| P<sub>β</sub> | – | – | 5.022(1) | 5.040(0) | 4.407(1) | 10.042(5) | 10.183(2) |

In the apo state (ssRNA bound NS3h), **MVIL** exhibits both ‘open’ and ‘closed’ conformations. In C1 (row 1, middle panel of Figure 6), the R461 and R464 side-chains are oriented inward, towards the ATP-pocket, and both residues are in close proximity to N417 (∼ 7 Å). Additionally, R464 is also near to H195 (∼ 7 Å). This inward orientation of R461 and R464 in C1 results in a ‘closed’ valve-like conformation of **MVIL** around the ATP-pocket and we hypothesise that this is a nucleotide stabilizing conformation at the ATP-pocket. Conversely, in C2 (row 1, right panel of Figure 6), R461 moves further inward towards the ATP-pocket, where it is surrounded by E286, N417, and Q457, with a separation distance of ∼ 4-5 Å. Notably, in C2, R464 undergoes an outward motion relative to the ATP-pocket, representing a significant change compared to C1. This outward movement causes R464 to shift from ∼ 21 Å away (in C1) to ∼ 4 Å (in C2) close to D471, forming a salt bridge between R464 and D471. These rearrangements make C2 as ‘open’ valve-like conformation. Our simulation started from C1 (‘closed’) conformations of **MVIL**; however, we noticed 37 transitions from ‘open’ to ‘closed’ in 5 *µ*s of simulation. This suggests R464 can spontaneously transition from the ‘open’ to ‘closed’ conformation within the limit of thermal energy, but the equilibrium is shifted towards the ‘open’ conformation due to large sampling of C2. It can be interpreted as the functional mechanism of nucleotide binding at the ATP-pocket: while **MVIL** is fluctuating between ‘open’ and ‘closed’ conformations, the nucleotide will encounter the ‘open’ conformation more frequently which facilitates nucleotide entry into the ATP-pocket. Following this, the **MVIL** transitions from the ‘open’ to ‘closed’ conformation where R464 interacts with the nucleotide to lock it into the ATP-pocket.

In the pre-hydrolysis (ssRNA+ATP-bound NS3h) state, the **MVIL** adopts ‘closed’ valve-like conformations to trap ATP at the ATP-pocket through an induction mechanism. Here, we observed the sampling of **MVIL** clusters C1 and C3. As suggested in the apo state, the inward motion of R461 and R464 in C1 (as in row 2, middle panel of Figure 6) forms close proximity to ATP to stabilize the nucleotide at the ATP-pocket. In this state, R461 is closer to the P*_γ_* (∼ 4 Å) than the P*_β_* (∼ 6 Å) of ATP. A similar trend is observed for R464, which is slightly closer to both the P*_γ_* (∼ 3.8 Å) and P*_β_* (∼ 5 Å) of ATP than R461. These subtle changes support the importance of C1 as a *selective* conformation for ATP stabilizer to NS3h, effectively acting as a ‘closed’ valve. However, the presence of ATP *induces* a structural rearrangement such as bringing E286, N417, and Q457 residues even closer ∼ 1-3 Å to R461 and R464 than in the apo state. Upon ATP binding, **MVIL** transitions to the C3 cluster (as in row 2, right panel of Figure 6), R461 moves closer to the P*_γ_* (∼ 3 Å) than in C1, but R464 maintains the same spatial distance from ATP phosphates. This results in N417 and Q457 moving ∼ 1 Å further away from R461 than in C1, while E286 moves closer to R461, though not as close as in C2. We hypothesize that these rearrangements in C3 lead the ATP-pocket (i.e. active site) to transition to a catalysis-active conformation while **MVIL** remains as a ‘closed’ valve by stabilizing the ADP portion of ATP through contact with R464.

In the post-hydrolysis-I (ssRNA+ADP+P_i_-bound NS3h) state, conformational induction of **MVIL** resists ADP release. The presence of both hydrolysis products *induces* sampling of the C4 cluster of **MVIL** in the ssRNA+ADP+P_i_ bound NS3h. In C4 (as in row 3, right panel of Figure 6), R461 moves ∼ 7 Å further from P_i_ as compared to P*_γ_* but maintains similar spatial separation from P*_β_* as in C1 and C3 of pre-hydrolysis state. This movement of R461 results in closer proximity to N417 and Q457 than C3. Similarly, R464 is also more distant from P_i_ as compare to P*_γ_*. Interestingly, R464 moves ∼ 1 Å closer to P*_β_* than in the pre-hydrolysis state. This results in the strongest affinity for ADP compared to the pre-hydrolysis state, thereby resisting ADP release.

In the post-hydrolysis-II (ssRNA+ADP-bound NS3h) state, **MVIL** adopts an ‘open’ valve-like conformation. This state, with only ADP present (or lacking Pi), leads to sampling of C1 and C6 MVIL loop conformations. In C1 (row 4, middle panel of Figure 6), R461 becomes ∼ 1 Å closer to P*_β_* than previous states. The motion of E413 and N417 surround the R461 is *selected* for C1 of apo state while closest proximity of E286 and Q457 to R461 is an *inductive* effect in the presence of (only) ADP. However, R464 moves ∼ 6 Å further away from P*_β_* but inward towards the ATP-pocket and becomes ∼ 5 Å closer to H195 than in the post-hydrolysis-I state. In the C5 cluster, R461 becomes subtly closer to P*_β_* (∼ 4.6 Å); more distant from N417 and Q457. In this cluster only, we observe R461 in close contact with E413 at ∼ 6.5 Å, as compared to the other clusters (∼ 10 Å). Most importantly, R464 separates from ADP, shifting away from the ADP plane (∼ 9 Å) or outward from the ATP-pocket than in C1 and becomes closest to H195 (∼ 4.5 Å). This opens the ATP-pocket, generating a repulsive effect on ADP and facilitating ADP release for the next hydrolysis cycle.

### Mutations at D471 Probe the Proposed Valve Behavior of MVIL

We propose D471 as a potential site for mutation to indirectly affect the ATP-pocket (i.e. active site) function. The R461 and R464 residues undergo critical inward motions during hydrolysis, causing the ATP-pocket to be ‘closed’, which is essential for ATP hydrolysis. Targeting these residues would disrupt the ATP hydrolysis function of WNV NS3h. However, this approach is common yet challenging for helicases due to specificity issues, as the active site motifs are widespread in host factors, potentially leading to detrimental effects. The outward orientation of R464 causing an ‘open’ ATP-pocket, likely affects ATP stabilization in the ATP-pocket (C2) or facilitates ADP release (C5). Both conformations involve the interaction of R464 with D471 or H195. While H194 is unsuitable for disruption due to its location in the P-loop, D471 is a viable choice because it is distant from the active site and solvent-exposed. This R464-D471 salt-bridge mediated **MVIL** ‘open’ valve-like conformation (C2) is only sampled in apo state along with ‘closed’ valve-like conformation (C1). Our proposal involves mutating D471 to maintain either a similar sidechain chemical nature or sidechain length through substitutions like D471E, D471N and D471L. We hypothesise that these mutation may favor any of the **MVIL** conformations observed in the Wild-Type apo state. This in turn, could impact ATP hydrolysis followed by replication inhibition by either destabilizing (‘open’) or blocking (‘closed’) of ATP binding.

To test this hypothesis, we generated D471 mutants in a WNV subgenomic replicon. This subgenomic replicon is a subviral genome capable of autonomous viral RNA replication but harboring a 5’ deletion of the structural genes C, prM, and E, which are replaced with a GFP reporter.^47^ This replicon cannot generate infectious virions but offers a sensitive approach to measure the impact of specific residues on viral RNA replication.We generated replicons with mutations at D471 (D471E, D471N, D471L) and performed qRT-PCR to assess effects on viral RNA replication. We found that substitution of D471 with either E or N reduced RNA replication 4-fold (Figure 7(a)). However, substitution of D471 with L reduced RNA replication 14-fold, to a level similar to a replicon harboring a mutation in the NS5 RdRp catalytic site, D668A (Figure 7A). We confirmed these data by immunoblotting for the GFP reporter, as well as the mutant NS3, from cells transfected with mutant replicons (Figure 7(b)). Additionally, we used automated fluorescent microscopy and quantified the number of cells expressing the GFP reporter for each mutant replicon (Figure 7(c) and Figure S3).

**Figure 7:**
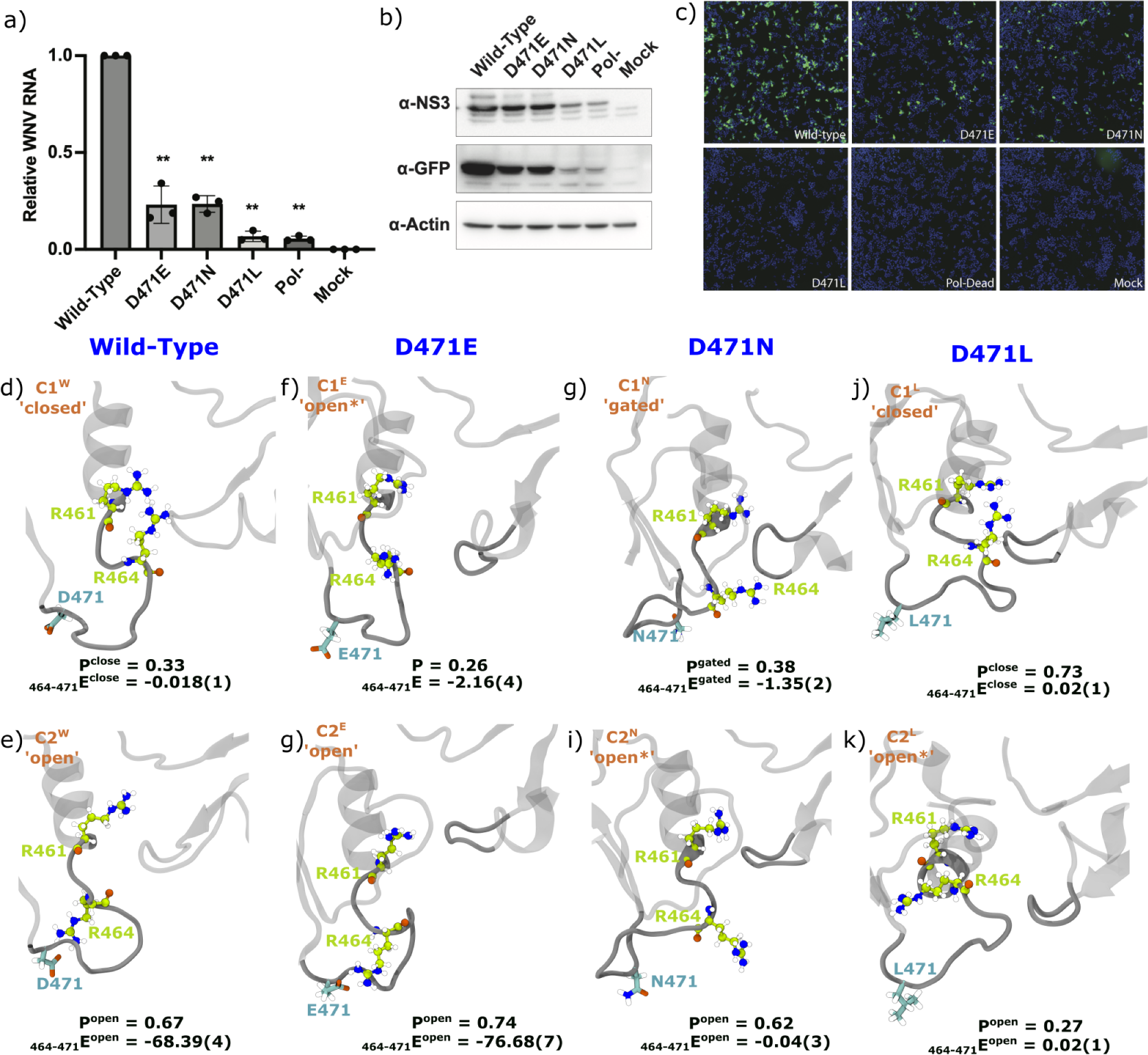
Mutations at D471 impact viral replication by influencing MVIL conformations to affect ATP binding. The top panel denotes experimental evidence for replication inhibition of mutants compared to the wild-type WNV: a) WNV replication assay of wild-type and D471 mutants (i.e. D471E, D471N and D471L), as well as a polymerase catalytic mutant Pol-, b) Western blot of cells transfected with mutant replicons, c) IF images of cells transfected with mutant replicons. GFP is whosh in green and nuclei are shown in blue. Data are represented as wild-type normalized, mean +/− SD, n=3. All statistics are one-way ANOVA with Dunnet’s correction for multiple comparisons **p<0.01 ***p<0.001. In the middle and lower panel, we present representative structures of **MVIL** clusters of wild-type (WT) and mutants identified by shapeGMM clustering: d-e) WT, f-g) D471E, h-i) D471N and j-k) D471L. Marking of each conformation is based on the 461-464 and 464-471 sidechain distances those are tabulated in Table S5. In each structural illustration, the probability (P) and interaction energy (_464–471_E in kcal.mol^−1^) between 464-471 is annotated for its respective **MVIL** clusters. For _464–471_E, the standard-error of last digit is presented in parentheses

To understand the mechanism of replication inhibition by the mutants, we simulated D471E, D471N and D471L WNV NS3h bound to ssRNA (apo state) followed by identifying unique conformational clusters of MVIL (heavy atoms of residues 461 to 472) with weighted ShapeGMM (W-SGMM). The scan analyses shows that each mutant explores two unique conformations of **MVIL** (see Figure S2). Due to differences in particles of **MVIL** between wild-type and mutants, we used sidechain distances of R461-R464 and R464-D471 to mark the clusters as ‘open’ or ‘closed’ (see Table S5). When R464 is inward and close to R461, we denote such cluster as ‘open’ while R464 is outward but forming salt-bridge with D471 that cluster as ‘closed’. With this description, Wild-Type C1^W^ and C2^W^ marked as ‘closed’ and ‘open’ respectively where the ratio of sampling is (0.33 : 0.67) (Figure 7(d-e)).

The **MVIL** of D471E destabilizes ATP binding for hydrolysis. We chose D471E mutation as a subtle modification which can maintain the chemical nature of wild-type while extending the sidechain. The **MVIL** explored two conformations C1^E^ and C2^E^ with sampling probability of 0.26 and 0.74 respectively (Figure 7(f-g)). We denote C1^E^ as ‘open*^∗^*’ conformation as R461-R464 are distant but R464 is not forming salt-bridge with E471 while R464-E471 forms salt-bridge in C2^E^ thus referred as ‘open’ conformation. Interestingly, sampling of **MVIL** ‘open’ conformation is increased in D471E compared to the wild-type which we believe is a manifestation of sidechain length extension. This indicates that D471E predominantly favors the ‘open’ ATP-pocket, allowing ATP to enter the active site. However, due to the strengthening of the R464-E471 salt-bridge compared to R464-D471, R464 is restricted from switching inward and is unable to stabilize (or lock) the ATP for hydrolysis, thereby reducing replication. Nevertheless, a smaller possibility of ATP stabilization exists due to the ‘open*^∗^*’ conformation, allowing ATP to enter and enabling R464 to move inward to bind ATP more strongly.

Next, we explored D471N mutation to evaluate the salt-bridge effect from maintaining the structure but changing the chemical nature from negatively charged to a polar residue. In this mutant also **MVIL** explored C1^N^ and C2^N^ clusters of conformations by sampling probability of 0.38 and 0.62 respectively (Figure 7(h-i)). In C1^N^, R464 is distant from both R461 and E471 but is closer to P-loop such that it is guarding the ATP-pocket, so we named this cluster as ‘gated’ which inhibits ATP to enter the active site. In contrast, R464 is distant from both R461 and E471 but not close to the P-loop, hence we designated this as ‘open’ and facilitating ATP entry. While C1^N^ represents an inhibitory conformation for ATP entry, its probability is smaller than the C2^N^. While we can not designate this as a predominant mode of replication reduction, this mutation underscores the importance of D471.

Mutating D471 to L blocks ATP entry to the active site. The motivation for ‘D’ to ‘L’ substitution is to maintain sidechain length but significantly change the chemical nature. However, D471L apo state only explored C1^L^ and C2^L^ clusters of **MVIL** with sampling probability of 0.73 and 0.27 respectively (Figure 7(j-k)). In C1^L^, R464 is inward and closure to R461 and thus we referring as ‘closed’ valve-like conformation. In contrast, R464 outward but unable to form contact with L471 and hence it marked as ‘open*^∗^*’ valve-like conformation. It suggest that D471L predominately favouring ‘closed’ valve-like conformation of **MVIL** in apo-state, opposite to the wild-type and blocks ATP entry followed by affecting ATP hydrolysis and thus replication reduction. However, existence of minor population of ‘open*^∗^*’ conformation of **MVIL** may help ATP binding which further induce inward motion of R464 to stabilize the ATP. Due to this possibility, a very small proportion of replication was observed.

## Conclusions

We presented our observations to support the differential binding enthalpy mechanism of WNV NS3h-ADP, which underscores hydrolysis turnover. We accomplished this by modeling hydrolysis states, including ssRNA, ssRNA+ATP, ssRNA+ADP+P_i_, and ssRNA+ADP-bound WNV NS3 helicase, and conducting separate microsecond-long equilibrium samplings of these states using all-atom, explicit water MD simulations. Due to the absence of substrate-bound WNV NS3h crystal structures, we employed the homology modeling method to create models, using the DENV structures ^20^ as templates. Our objective of this study is to provide molecular insights into protein and nucleotide interactions, and their relationship with nucleotide-dependent protein conformational changes. However, this force is not the only factor at play at the interface. Interactions with water also play a crucial role in the binding enthalpy. Simulations of DENV NS3h revealed a significant decrease in water molecules at the ATP-pocket in the pre-hydrolysis (ssRNA+ATP) state compared to the apo (ssRNA) state.^23^ This suggests that water has a more entropic effect on nucleotide binding. Our analysis does not consider the contribution of divalent cations (Mg^2+^). In ZIKV NS3h, Cao et al. reported the influence of divalent cations on changing the bound ATP conformation, along with motif-I, leading to a pre-hydrolysis state.^59^ Additionally, our calculations do not account for interactions with the P*_γ_* or P_i_ moiety; however, our aim is to conduct a comparative analysis with comparable parameters.

Despite the aforementioned limitations, our calculations revealed a significantly different and clear trend in the unique protein-nucleotide binding enthalpy for the hydrolysis states. A favorable enthalpic change between the post-hydrolysis-I and pre-hydrolysis states dictates NS3h’s progression towards hydrolysis. This altered binding enthalpy calculated for modeled WNV NS3h ensembles was not deducible from TBEV NS3h crystal structures,^30^ although these observations may be specific to the protein source (i.e., the virus). Interestingly, the unfavorable change in binding enthalpy between the post-hydrolysis-II and post-hydrolysis-I states suggests a dependence on the release of P_i_. Furthermore, evidence of the progression to the next hydrolysis cycle is also apparent through the favorable enthalpic change between the pre-hydrolysis and post-hydrolysis-II states. In contrast, Perez-Villa et al. report that ADP binds more strongly than ATP in HCV NS3h simulations.^29^ However, the authors suggest the possibility of an entropic effect at play. For a deeper molecular understanding, the breakdown of binding enthalpy into residue-level contributions revealed that the **MVIL** residues play a significant role. Most importantly, the affinity of R461 and R464 for ADP changes drastically among the hydrolysis states, and the motion of these residues is locally correlated within the **MVIL**, which covers the ATP-pocket (active site). Nucleotide-specific motion of R463 and R466 (R461 and R464 in WNV respectively) is also evident in the crystallographic report of TBEV.^30^

The **MVIL** structure adjacent to the active site (i.e., ATP-pocket) plays a significant role in ATP hydrolysis cycle through nucleotide-dependent loop conformations. The presence of a substrate (i.e. nucleotide) alters the conformational equilibrium of loops, which can be attributed to either a *selection* or *induction* mechanism, i.e., the existence of certain conformations or the emergence of new ones, respectively. In WNV NS3h, the **MVIL** exhibits a significant effect on altering the binding enthalpy for hydrolysis states, primarily due to the nucleotide-specific sampling of **MVIL** conformations. Our Shape-GMM based clustering and LDA analysis identified five unique clusters of MVIL. The computation of nucleotide-dependent sampling of **MVIL** cluster populations demonstrates a differential affinity of NS3h-ADP, predominantly driven by the *induction* mechanism along the hydrolysis cycle and to a smaller extent by the *selection* mechanism. The analyses suggest that stabilization of ATP after binding is a *selection* process, and subsequently, the presence of ATP *induces* MVIL into a new cluster, which may represent the catalytic active state. After hydrolysis, the presence of products further *induces* conformational changes in **MVIL**, strongly grasping the ADP. The presence (absence) of ADP (P_i_) in the post-hydrolysis-II state leads to both conformational *selection* and *induction* in MVIL, with a similar trend observed in the apo state, possibly reflecting the *induction* of ADP release.

Comparing the **MVIL** clusters at the molecular level revealed that the differential motion of R461 and R464 leads **MVIL** to function as a dynamic valve for hydrolysis states. The inward motion of both R461 and R464 residues towards the ATP-pocket causes C1, C3, and C4 **MVIL** conformations to act as a ‘closed’ valve for the ATP-pocket. The spatial arrangement of R461, R464, and surrounding residues contributes to the formation of a strong affinity for ADP in the pre-hydrolysis and post-hydrolysis-I states. However, in C4, R464 displays a subtle closure to ADP compared to other clusters, resulting in the highest NS3h-ADP affinity in the post-hydrolysis-I state. Importantly, during our simulation, we observed the sampling of C2 and C5 clusters, which represent an ‘open’ valve-like structure of **MVIL** due to the outward motion of R464 in the apo and post-hydrolysis-II states respectively. It is worth noting that our simulation began with the inward orientation of R464 in each hydrolysis state. In C2, the outward motion of R464 leads to the formation of a new salt-bridge with D471. This is the first report of the possibility of an R464-D471 salt-bridge, which results in an ‘open’ valve-like structure of **MVIL** in the apo state (NS3h bound to ssRNA). Moreover, D471 is a conserved residue within the *Flavivirus* genus. Thus, it indicates this salt-bridge formation could be a common structural feature and a potential site of indirect disruption of active site function. However, in C5, the outward motion of R464 represents a kind of register-shift motion compared to the ADP plane, resulting in the weakest affinity for ADP in the post-hydrolysis-II state. This arrangement causes **MVIL** to act as an open valve for product release. This detachment trend of R464 from ADP in the post-hydrolysis-II state is also reported by Anindita et al. in the TBEV NS3h crystal structure.^30^

Our clustering analyses of **MVIL** and R464 motion motivated us to mutate D471 to indirectly disrupt hydrolysis at the ATP-pocket and assess impacts on viral replication. Our western blot results and replication assays showed that, while we observe NS3h protein expression from the D471E, D471N and D471L replicon constructs, these mutations significantly reduce WNV replication. Modelling and simulation of these mutants in the apo state show sampling of two conformational clusters of **MVIL**. The D471E and D471L mutation display shifts from the wild-type population of **MVIL**, predominantly sampling ‘open’ and ‘closed’ valve-like conformations, respectively. These observations suggest a probable mechanism of replication reduction: D471E is unable to stabilize ATP in the ATP-pocket after ATP binds in the ‘open’ conformation, while in D471L NS3h, ATP is unable to enter the ATP pocket due to a ‘closed’ ATP-pocket. Interestingly, in the D471N simulation we could not discern a probable mechanism of hydrolysis disruption, but observed sampling of closed-like conformations of **MVIL** to a smaller extent which can impact ATP entry. Altogether, these mutations highlight the importance of D471, serving as a potential site to alter vale-like conformational dynamics of **MVIL**, indirectly targeting the ATP-pocket of a critical viral enzyme.

Our exhaustive simulations, rigorous analyses and experimental evidence have showcased the differential affinity-based functional mechanism of NS3h hydrolysis turnover, regulated by the valve-like dynamics of the **MVIL** conformations. Analogous to a combustion engine, initially (apo state), MVIL exists as an open valve. Once the fuel (i.e., ATP) enters the chamber (i.e., ATP-pocket), **MVIL** transforms into a closed valve, locking the fuel (prehydrolysis and post-hydrolysis-I states). After combustion (i.e., ATP hydrolysis), the valve opens the chamber for the release of the product (i.e., ADP in post-hydrolysis-II state). However, this is not a simple mechanical process but is underlined by the complex dynamics of conformational induction. This study provides an in-depth understanding of the critical role of the **MVIL** in hydrolysis states in WNV NS3h, beyond its simple function as a nucleotide stabilizer. The **motif VI loop** is a surface-exposed structure and can be leveraged to target the hydrolysis state-dependent conformation to abolish its hydrolysis function as observed in our D471 mutation rather than directly targeting the ATP-pocket, which could be detrimental to host helicases. Structurally, NS3h shares a similar arrangement in the *Flavivirus* genus, and we believe these insights will aid in understanding other virus helicases. In addition to catalysis, the active site loop possesses a triggering or triggered loop effect, as seen in the WPD loop.^32^ Future work in our group will explore such possibilities in the WNV NS3h helicase.

## Supplementary data

Supplementary data includes four tables (Table S1 to S5) and two figures (Figure S1 to S3).

## Acknowledgement

The funding for this project was supported by the National Institute of Health (NIH) NIAID Award R01AI166050 (MM), RO1AI143850 (HR) and NIH NIGMS Award T32GM144302 (ZW). The computing for this project was performed at the High Performance Computing Center at Oklahoma State University supported in part through the National Science Foundation Grant OAC-1531128. We also acknowledge Anvil Supercomputer through ACCESS to provide the computing hours for our analyses.

## TOC Graphic

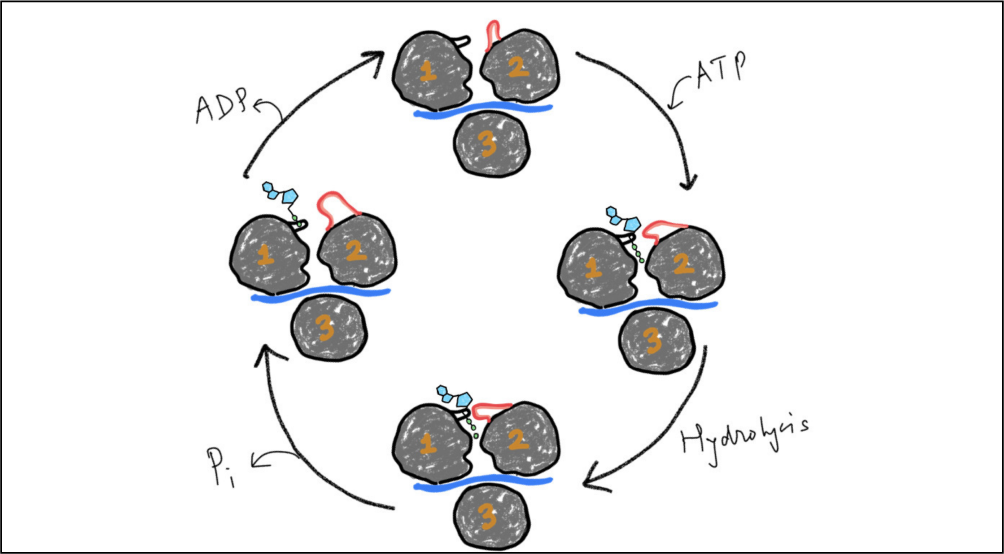

