## Supporting Information for "Motif-VI Loop Acts as a Nucleotide Valve in the West Nile Virus NS3 Helicase"

Table S1: **Maintenance of octahedral coordination of $Mg^{2+}$ within 2.5 Å for hydrolysis substrate and products bound WNV NS3h+ssRNA.**

| **Atom Name** | **ATP** | **ssRNA+ATP** | **ssRNA+ADP+$P_i$** | **ssRNA+ADP** |
|---|---|---|---|---|
| $OG1_{T200}$ | 99.94% | 99.98% | 99.63% | 99.84% |
| $OE2_{E285}$ | 99.94% | – | – | – |
| $O3G_{ATP}$ | 100% | – | – | – |
| $O2G_{ATP}$ | – | 100% | – | – |
| $O1B_{ATP}$ | 100% | 100% | – | – |
| $O1B_{ADP}$ | – | – | 100% | 100% |
| $O3_{P_i}$ | – | – | 100% | – |
| $O_{W1}$ | 100% | 100% | 100% | 100% |
| $O_{W2}$ | 100% | 100% | 100% | 100% |
| $O_{W3}$ | – | 100% | 100% | 100% |
| $O_{W4}$ | – | – | – | 100% |

Table S2: **Occurrence probability of conserved contacts in different substrates bound WNV NS3 helicase models with ssRNA.** A 5 Å of cut-off distance is used between atoms of protein and ssRNA. The residue identity in WNV NS3h is similar to DENV.

| **Motif** | **ZIKV** | **DENV** | **ssRNA** | **ssRNA+ATP** | **ssRNA+ADP+$P_i$** | **ssRNA+ADP** |
|---|---|---|---|---|---|---|
| **Ia** | P224 | P223 | 99% | 99% | 99% | 99% |
| **Ia** | T225 | T224 | 99% | 81% | 99% | 99% |
| **Ia** | R226 | R225 | 99% | 99% | 99% | 99% |
| **Ia** | – | V226 | 99% | 99% | 99% | 99% |
| **II** | T290 | T289 | 64% | 8% | 6% | 63% |
| **II** | D291 | D290 | 99% | 99% | 99% | 99% |
| **IV** | P364 | P363 | 7% | 93% | 8% | 99% |
| **IV** | S365 | S364 | 35% | 99% | 97% | 99% |
| **IV** | V366 | I365 | 12% | 98% | 99% | 99% |
| **IV** | K367 | R366 | 5% | 56% | 84% | 95% |
| **IVa** | S387 | S386 | 2% | 99% | 99% | 9% |
| **IVa** | R388 | R387 | 99% | 99% | 99% | 99% |
| **IVa** | K389 | K388 | 4% | 63% | 93% | 8% |
| **V** | T409 | T408 | 36% | 99% | 99% | 98% |
| **V** | D410 | D409 | 97% | 99% | 99% | 4% |
| **V** | I411 | I410 | 52% | 99% | 99% | 98% |

Table S3: **Occurrence probability of conserved contacts formed by WNV-NS3h with ATP, ADP+$P_i$ and ADP in presence of ssRNA.** In ZIKV and DENV, the contacts are identified from crystal structure. In ZIKV, only ATP+ssRNA bound NS3h is considered (PDB:7V2Z), while in DENV all hudrolysis substrate and products bound structures (PDB:2JLV, 2JLY, 2JLZ) are taken into account. A 5 Å of cut-off distance is used between atoms of protein and ATP or ADP or $P_i$.

| **Motif** | **ZIKV** | **DENV** | **ssRNA** | **ssRNA+ATP** | **ssRNA+ADP+$P_i$** | **ssRNA+ADP** |
|---|---|---|---|---|---|---|
| **I** | H195 | H194 | H194 | 61% | 86% | 3% |
| **I** | P196 | P195 | P195 | 99% | 99% | 97% |
| **I** | G197 | G196 | G196 | 99% | 99% | 99% |
| **I** | A198 | A197 | A197 | 99% | 99% | 99% |
| **I** | G199 | G198 | G198 | 99% | 99% | 99% |
| **I** | K200 | K199 | K199 | 99% | 99% | 99% |
| **I** | T201 | T200 | T200 | 99% | 99% | 99% |
| **I** | – | K201 | R201 | 99% | 67% | 84% |
| **I** | – | R202 | R202 | 97% | 9% | 49% |
| **Ia** | – | E230 | E230 | 5% | 32% | 27% |
| **II** | E286 | E285 | E285 | 63% | 98% | 68% |
| **III** | – | A316 | A316 | 8% | 75% | – |
| **V** | M414 | M413 | M413 | 1% | 99% | 28% |
| **V** | G415 | G414 | G414 | 20% | 99% | 23% |
| **V** | – | A415 | A415 | 88% | 18% | 56% |
| **V** | N417 | N416 | N416 | 92% | 90% | 94% |
| **VI** | Q455 | Q456 | Q456 | 8% | 98% | – |
| **VI** | R459 | R460 | R460 | 99% | 99% | 97% |
| **VI** | R462 | R463 | R463 | 99% | 99% | 70% |
| **VI** | N463 | N464 | N464 | 5% | 99% | 97% |

Figure S1: **Linear correlation of protein residues with R460 and R463.** Presented correlation is the Pearson's correlation of $C_\alpha$ atoms of protein. Averaged over the entire trajectory.

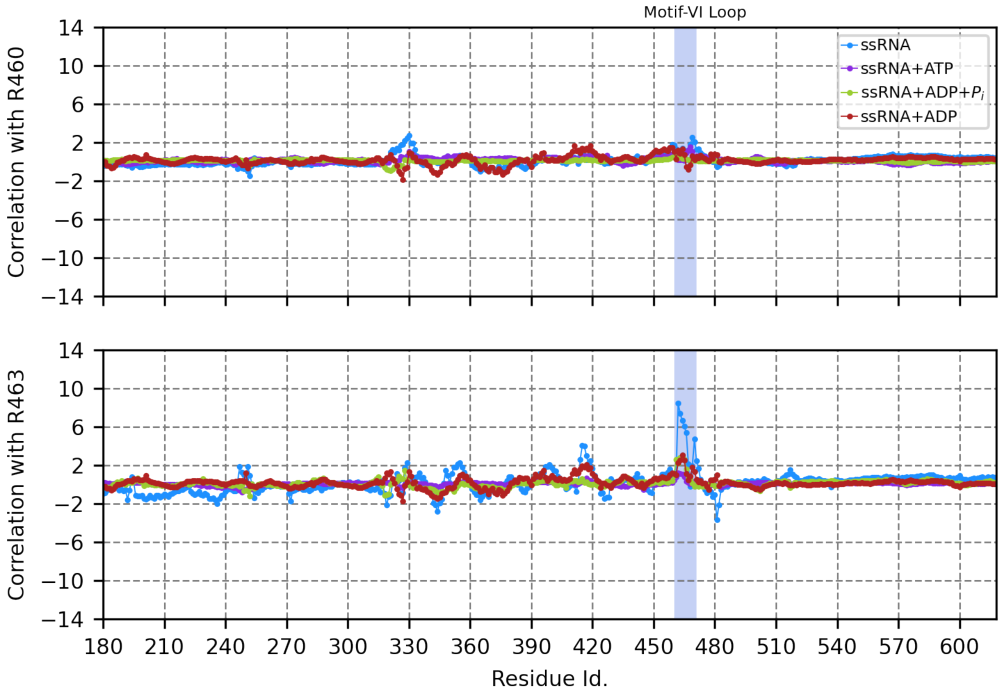

Table S4: **Significance of $E_{inter}^{NS3h-ADP}$ distributions with $t$-test.** We randomly selected 501 dataset from each of the two compared distributions. Same step of new random dataset is iterated for 1000K times. The reported absolute $t$-value is averaged over the iterations which is higher than the critical value (1.96) for 0.05 significance level of 1000 degrees of freedom (df)

| **Compairing Systems** | ***t*-value** |
|---|---|
| ssRNA+ATP and ssRNA+ADP+$P_i$ | 44.02(1.45) |
| ssRNA+ADP+$P_i$ and ssRNA+ADP | 77.61(2.0) |
| ssRNA+ATP and ssRNA+ADP | 45.11(1.5) |

Figure S2: **Clustering of motif-VI loop ensembles sampled in the mutants of WNV NS3h apo state.** The log likelihood per frame as a function of clusters is plotted for each mutant with change of slope measure by $2^{nd}$ derivative. The clustering is performed with ShapeGMM algorithm and the feature consists with positions of all heavy atoms of residues 461 to 472. Each training run is iterated for 5 times for each cluster.

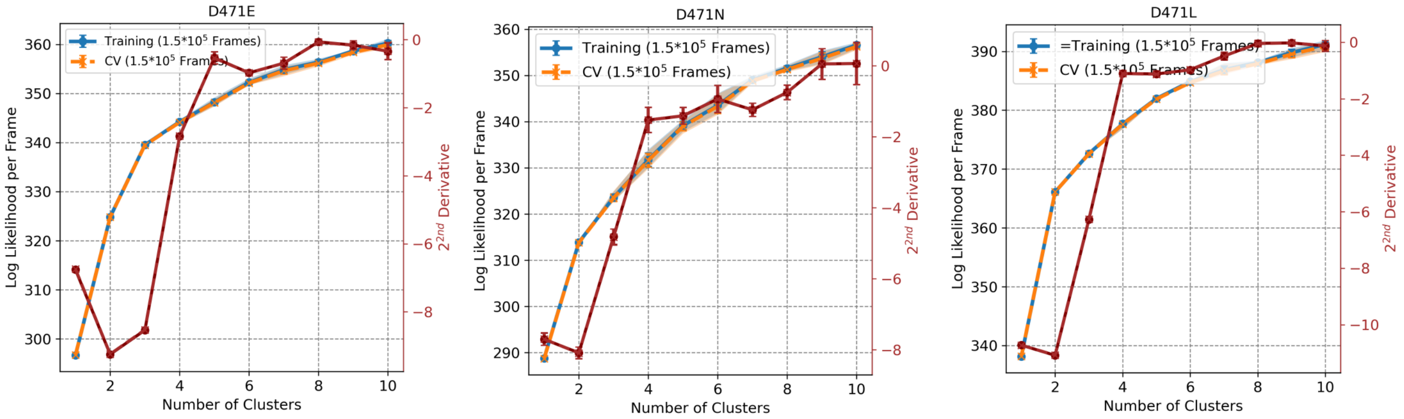

Table S5: **Cluster specific distances between key residues of MVIL.** To define the loop **MVIL** loop stattus we used sidechain distances of 461-464 and 464-471 residue pair. We chose terminal sidechain 'C' position of any residue to compute the distance. Parentheses vale denote standard error of last digit. Units is $\AA$. See text for valve status definition details.

| **Residue Pair** | **WT** | | **D471E** | | **D471N** | | **D471L** | |
|---|---|---|---|---|---|---|---|---|
| | $C1^W$ | $C2^W$ | $C1^E$ | $C2^E$ | $C1^N$ | $C2^N$ | $C1^L$ | $C2^L$ |
| 461-464 | 5.056(4) | 18.060(2) | 12.33(1) | 16.746(2) | 12.547(7) | 13.079(4) | 7.814(3) | 9.832(7) |
| 464-471 | 19.456(3) | 4.837(2) | 16.41(1) | 5.148(5) | 12.63(1) | 16.179(6) | 20.722(3) | 20.27(1) |
| Valve Status | **closed** | **open** | **open**$^*$ | **open** | **gated** | **open**$^*$ | **closed** | **open**$^*$ |

Figure S3: **Quantification of reporter GFP expression for each replication construct.** Data are represented as wild-type normalized, mean +/- SD, n=3. All statistics are one-way ANOVA with Dunnet's correction for multiple comparisons **p<0.01 ***p<0.001.

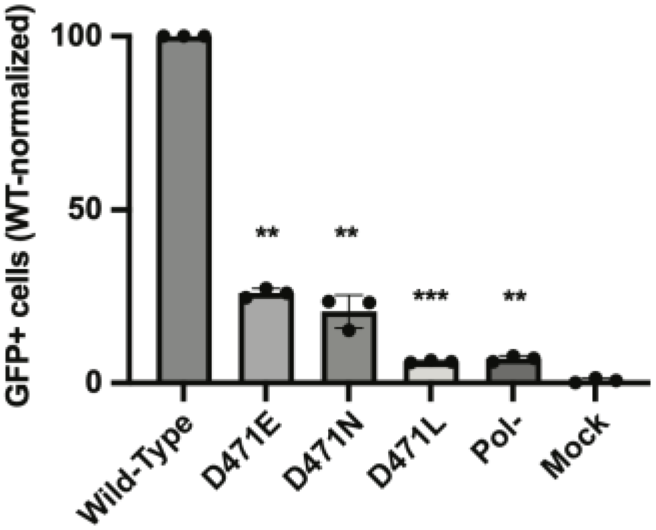
